# Endogenous cAMP production within tissue fibroblasts is required for spontaneous fibrosis resolution

**DOI:** 10.64898/2026.11.12.765422

**Authors:** Sean M. Fortier, Jennifer Speth, Natalie M. Walker, Jared D. Baas, Kevin Wang, Vivian T. Wong, Steve D. Guzman, Yang Liu, Haichun Pan, Hao Ren, Priyanka Verma, Steven K. Huang, Sergey Gutor, Timothy S. Blackwell, John Varga, Carlos A. Aguilar, Rachel L. Zemans, Marc Peters-Golden

**Author notes:** <u>Corresponding Author</u>: Sean M. Fortier, MD, 6301B MSRB III, 1150 W. Medical Center Dr. Ann Arbor, MI 48109-5642.

## Abstract

Pathologic myofibroblasts (MFs) are pivotal effector cells in the development of persistent tissue fibrosis following parenchymal injury, and their activation and accumulation has been linked to a failure to phenotypically dedifferentiate and thus to resist apoptosis. The second messenger cyclic AMP (cAMP) has been shown to induce MF dedifferentiation and restore their sensitivity to apoptosis *in vitro*, but its endogenous role within MFs during fibrosis resolution *in vivo* has not been explored. To test the hypothesis that cAMP generation and signaling within fibroblasts is essential for fibrosis resolution, we leveraged the spontaneously resolving nature of the single-dose intrapulmonary and subcutaneous bleomycin models in mice with an inducible and fibroblast-specific abrogation of cAMP generation owing to deletion of the stimulatory G protein (Gαs). Abolishing cAMP generation in fibroblasts after peak lung fibrosis was established led to a failure of the resolution with single-cell RNA seq demonstrating conspicuous differences in pathway enrichment and cellular cross-talk not only in fibroblasts, but also within macrophages and epithelial cells. Spontaneous resolution of skin fibrosis in a model of scleroderma similarly required cAMP production in dermal fibroblasts. We conclude that cAMP generation within fibroblasts after tissue fibrosis has been established is required for its spontaneous resolution in the lung and skin. Our findings provide new insights into factors governing the outcome of tissue fibrosis as well as a mechanistic basis that helps to explain the therapeutic efficacy and ultimate potential of newly available and emerging cAMP-elevating agents.

## INTRODUCTION

Tissue fibrosis is a final common pathway accounting for organ dysfunction across a wide range of chronic diseases (*1, 2*). Scarring of the lungs is especially devastating as it leads to respiratory failure from parenchymal stiffening, distortion of tissue architecture, and impaired alveolar gas exchange (*3*). The hallmark pathobiology of progressive fibrotic lung diseases such as idiopathic pulmonary fibrosis (IPF) and systemic sclerosis/scleroderma is characterized by defective alveolar epithelial regeneration and repair, dysregulated cellular crosstalk, and a persistence of pathological and aberrant cell types (*4*). Among these cells is the myofibroblast (MF), traditionally considered the ultimate effector of tissue fibrosis. Following lung injury resident fibroblasts undergo phenotypic differentiation to MFs, which produce excess collagens and other ECM proteins, and transmit tensile forces through stress fibers and focal adhesions (*5*). In contrast to their clearance through phenotypic dedifferentiation and apoptosis during normal repair, persistent parenchymal scarring is associated with the accumulation of apoptosis-resistant MFs (*6, 7*). Understanding the operative molecular pathways that dictate whether injured lungs undergo resolution or become persistently fibrotic is fundamental to our capacity to intervene therapeutically. However, the endogenous events essential to mediating MF dedifferentiation – a demonstrated determinant of fibrosis resolution (*8*) – have not been elucidated.

Certain animal models of pulmonary fibrosis are characterized by spontaneous resolution (*9, 10*). Perhaps the most well-known and frequently employed is the single-dose bleomycin (bleo) murine model. Following intrapulmonary bleo administration in healthy young mice, damage to lung epithelium prompts inflammation and injury through the first 6-9 days followed by a fibrotic phase involving fibroblast accumulation and excess collagen deposition that peaks at day 21 and naturally resolves over the ensuing 6 weeks (*11*). Though often framed as a critical and unfortunate limitation, this self-resolving nature of the bleo model provides an opportunity to interrogate endogenous pathways that favor resolution over persistent fibrosis. Utilizing genetically modified mice containing an inducible site-specific recombinase, putative anti-fibrotic brakes can be deleted in cells of interest to determine if they are in fact necessary to resolve fibrosis. This approach has been used to demonstrate that Fas-mediated apoptosis of lung fibroblast populations is indeed necessary to permit spontaneous fibrosis resolution (*12*).

We have previously proposed that MF dedifferentiation may serve as a surrogate readout to identify pathways and agents capable of promoting resolution of fibrosis in animal models (*13*). Indeed, an extensive body of literature attests to the ability of the cAMP axis to not only robustly inhibit nearly all activation functions of lung fibroblasts (*14–16*), but to durably reverse the phenotype of differentiated lung MFs, as reflected by global remodeling of their transcriptome (*17, 18*). Moreover, tissue fibrosis has been associated with a myriad of disruptions in the cAMP signaling axis (*19*), suggesting its central role as an endogenous determinant of lung homeostasis. Akin to the requisite inactivation of tumor suppressors enabling malignant transformation in cancer, disabling cAMP generation in lung fibroblasts may be required for fibrosis to progress and persist. Longitudinal single cell transcriptomic analysis of mouse lungs during spontaneous resolution reveals enrichment of cAMP pathways within lung fibroblasts, macrophages, and epithelial cells (*20*). Nevertheless, the necessity of endogenous cAMP production and signaling within lung fibroblasts to promote fibrosis resolution has not been demonstrated and its specific role during resolution has not been characterized.

With rare exceptions, intracellular cAMP generation in mammalian cells depends on ligation of a G protein-coupled receptor (GPCR) that activates adenylyl cyclase (AC) (*21*). Given the multitude of ligand-GPCR pairs capable of increasing cAMP levels in lung fibroblasts, deleting any single receptor would be insufficient to abolish its production. However, as all GPCRs coupled to cAMP generation require Gαs (encoded by Guanine Nucleotide-Binding Protein, Alpha Stimulating gene: *GNAS*), we sought to exploit this physiologic bottleneck to investigate the role of cAMP signaling during fibrosis resolution. Using CRISPR/Cas9 and loxP Cre-mediated deletion, we verified that *GNAS* was necessary for *in vitro* cAMP production and MF dedifferentiation elicited by the well-characterized antifibrotic prostanoid prostaglandin E_2_ (PGE_2_). Moreover, *in vivo* fibroblast-specific *Gnas* deletion induced after the development of peak bleo-induced fibrosis abrogated spontaneous resolution as observed in WT mice and instead led to persistent fibrosis and failed/dysregulated alveolar repair. Single cell RNA-seq (scRNA-seq) revealed conspicuous differences in pathway enrichment and cellular crosstalk among mesenchymal, epithelial, and immune cell populations in resolved (WT) versus fibrotic (fibroblast-specific *Gnas*-deficient) lungs. Mirroring these effects in the lung, cAMP signaling in dermal fibroblasts was likewise necessary for spontaneous fibrosis resolution in the skin in a mouse model of scleroderma employing subcutaneous bleo administration. These findings establish cAMP as a critical endogenous mediator whose timely production within fibroblasts following tissue injury is required to orchestrate spontaneous fibrosis resolution. They also help to explicate the mechanisms of action of two new therapeutic agents for pulmonary fibrosis, nerandomilast and treprostinil.

## RESULTS

### *Gnas* deletion abrogates GPCR-mediated cAMP generation and enhances pro-fibrotic gene expression and apoptosis resistance

To further corroborate and complement existing literature (*19*) indicating that cAMP signaling is impaired within fibroblasts in fibrotic states, we determined levels of phosphorylated cAMP response-element binding protein (P-CREB) – a transcription factor activated by cAMP/PKA-mediated phosphorylation – in cells from normal and fibrotic human and mouse lungs. P-CREB was reduced in IPF as compared to normal fibroblasts as determined both by Western blot analysis of cultured cells (Supplemental Figure 1A) and *in situ* immunofluorescence staining (Supplemental Figure 1B). As expected, fibroblasts within lung sections from IPF patients concomitantly demonstrated substantially higher α-smooth muscle actin (αSMA) expression than did cells from normal lungs. A similar pattern of reduced cAMP signaling as judged by P-CREB staining within tdTomato-labeled fibroblasts was also observed in mouse lungs at peak fibrosis following bleo administration (Supplemental Figure 1C).

We next sought to phenocopy this impairment in cAMP signaling observed in fibrotic fibroblasts using CRISPR/Cas9-mediated knockdown of the *GNAS* gene encoding Gαs in human lung fibroblasts (Figure 1A left). In unstimulated fibroblasts in culture, cAMP levels were essentially undetectable (Figure 1A right), which is consistent with the short half-life of intracellular cAMP owing to the action of cAMP-degrading phosphodiesterases. We therefore treated fibroblasts with prostaglandin E_2_ (PGE_2_), which ligates two Gαs-coupled GPCRs (EP2 and EP4), and measured cAMP concentrations in stable lines expressing Cas9 and either non-targeting (NT) or *GNAS*-specific guide RNAs (sgRNA). Fibroblasts harboring a non-targeting (NT) sgRNA exhibited robust increases in cAMP 5 min following exposure to PGE_2_. In contrast, Gαs null fibroblasts treated with PGE_2_ generated significantly lower levels of cAMP (Figure 1A right, bottom). The failure to completely abolish cAMP production is likely a consequence of incomplete CRISPR/Cas9 deletion as indicated by Western blot (Figure 1A right, top). Nevertheless, *GNAS* KO human lung fibroblasts maintained in culture demonstrated higher baseline levels of collagen I and αSMA (Figure 1B) and this expression correlated with a relative resistance to anti-Fas antibody-mediated apoptosis sensitivity in MFs as measured by annexin V expression (Figure 1C). These results suggest the presence of a baseline Gαs-dependent inhibitory cAMP tone that limits fibrosis-associated genes and increases apoptosis sensitivity in culture. Finally and importantly, Gαs-deficient MFs failed to undergo dedifferentiation (as reflected by collagen I expression) following treatment with PGE_2_, although they did so following treatment with the cell-permeable cAMP analog dibutyryl cAMP, which bypasses the need for GPCR-coupled cAMP generation (Figure 1D).

**Figure 1.**
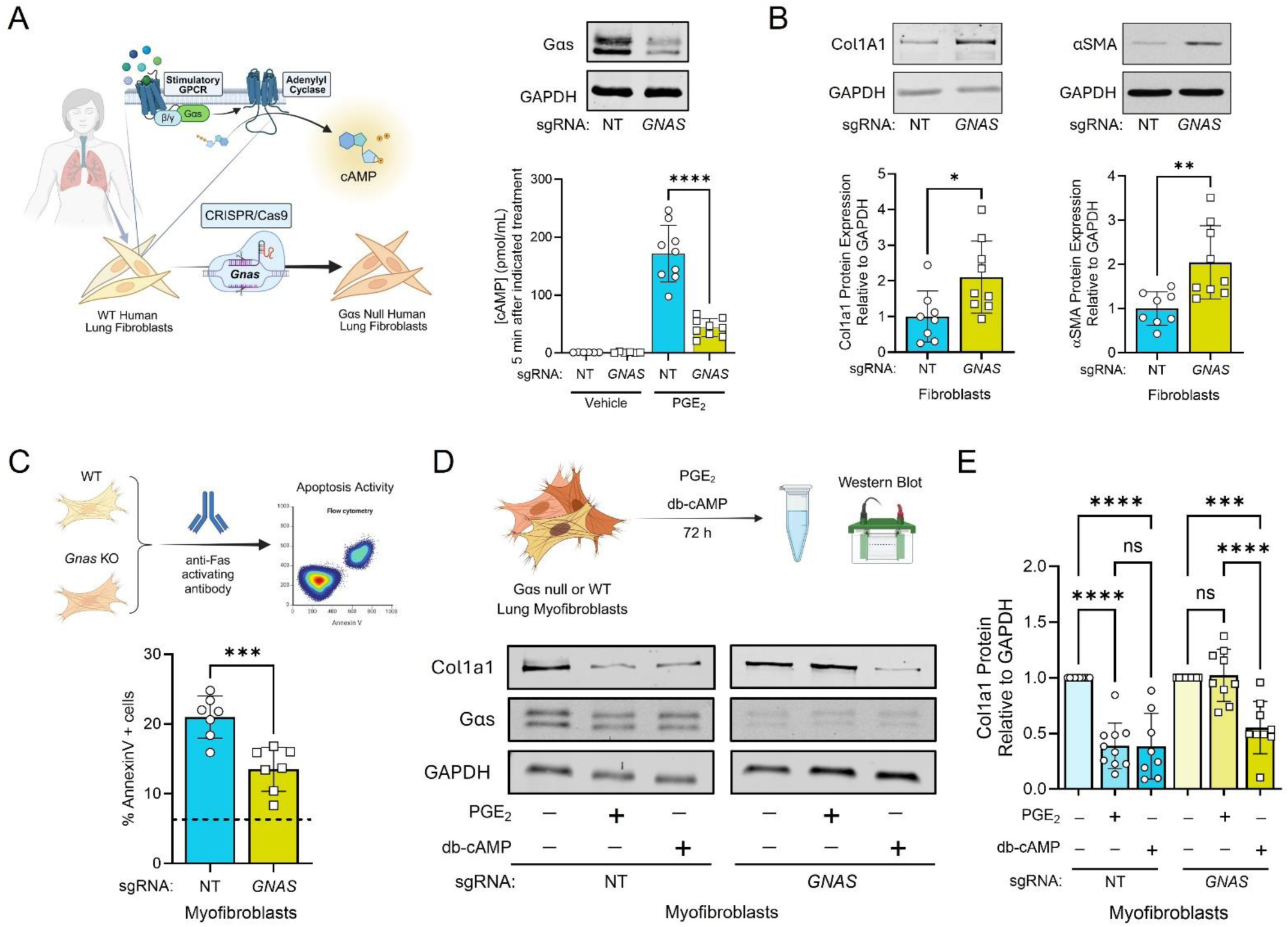
*GNAS* deletion abrogates GPCR-mediated cAMP generation and enhances baseline pro-fibrotic gene expression and apoptosis resistance in lung fibroblasts. (**A**) CRISPR/Cas9-mediated Gαs deletion using *GNAS* or non-targeting (NT) sgRNAs in MRC5 fibroblasts and its impact on PGE_2_-stimulated cAMP generation. (**B**) Protein quantification by Western blot of fibrosis-associated genes Col1A1 and αSMA in unstimulated MRC5 fibroblasts treated with *GNAS* or NT sgRNAs and maintained in culture with serum free medium for 48 h (top: representative blots, bottom: densitometric analysis). (**C**) Myofibroblast apoptosis sensitivity as measured by % Annexin V staining. Cultured WT and *GNAS* KO (via CRISPR/Cas9 deletion) MRC5 fibroblasts following exposure to TGFβ (2 ng/mL) for 48 h were subsequently treated with anti-Fas activating antibody (100 ng/mL) for 24 h followed by harvest for flow cytometry. Dashed line indicates % Annexin V+ cells in WT fibroblasts not treated with anti-Fas activating antibody. (**D**) WT and Gαs null MRC5 fibroblasts were treated with TGFβ (2 ng/mL) for 48 h to generate MFs (top). Collagen 1A1 expression was quantified in MFs treated with either PGE_2_ (500 nM) or db-cAMP (2 mM) for 72 h (bottom). (**E**) Densitometric analysis of collagen expression in D. In all panels, the sample number for each experiment (n) varied between 8 and 10 and is indicated by the number of data points in each histogram. \**P <* 0.05, ** *P <* 0.01, \*\*\**P <* 0.001, and \*\*\*\**P <* 0.0001 were determined by 2-tailed *t* test (**A** – **C**) and 1-way ANOVA (**E**).

### Endogenous cAMP production within lung fibroblasts promotes their apoptotic clearance and is necessary for spontaneous fibrosis resolution

Our *in vitro* human data indicate that Gαs knockdown (via *GNAS* deletion) effectively reduces GPCR-mediated cAMP production in lung fibroblasts. We therefore reasoned that its fibroblast-specific deletion *in vivo* can be leveraged to determine the role of endogenous fibroblast cAMP signaling during spontaneous fibrosis resolution. We established a *Col1a2^CreERT+/0^;Gnas^fl/fl^* mouse enabling tamoxifen-induced deletion of *Gnas* in fibroblasts (herein referred to as *Col1a2;Gnas* KO fibroblasts). We first tested whether this Cre-mediated *Gnas* deletion model successfully abrogated cAMP generation in mouse lung fibroblasts. Fibroblasts obtained from the lungs of *Col1a2^CreERT+/0^;Gnas^fl/fl^* mice maintained on a tamoxifen chow diet for 2 weeks demonstrated markedly diminished cAMP production in response to PGE_2_ compared to WT lung fibroblasts (Supplemental Figure 2A) similar to the effect of CRISPR/Cas9-mediated *GNAS* deletion in human lung fibroblasts (Figure 1A) and with analogous effects on pro-fibrotic gene expression (Supplemental Figure 2B).

For subsequent *in vivo* experiments, we incorporated a Cre-inducible tdTomato fluorescent tag (*R26^tdT+/0^*) into our mouse line to enable facile identification of fibroblasts. To evaluate the necessity of cAMP signaling within lung fibroblasts for spontaneous fibrosis resolution, we employed the single-dose bleo model. Specifically, 1.0 U/kg bleo or an equal volume of saline was administered by oropharyngeal (o.p.) aspiration to *Col1a2^CreERT+/0^;R26^tdT+/0^* control mice, which were harvested on day 21 to assess peak fibrosis. In this group, a tamoxifen chow diet was introduced on day 6 through 21 to induce tdTomato expression. In parallel, 1.0 U/kg bleo was similarly administered to *Col1a2^CreERT+/0^;Gnas^fl/fl^;R26^tdT+/0^* or *Col1a2^CreERT+/0^;R26^tdT+/0^* controls. Mice in this cohort were maintained on a tamoxifen chow diet starting on day 21 through harvest on days 42 or 63 to assess mid- and full resolution (*22*), respectively (Figure 2A). Deletion of *Gnas* after peak fibrosis served to ensure that outcomes can be attributed to the influence of fibroblast-specific cAMP signaling exclusively during the resolution process. Successful Cre-mediated recombination of the *Gnas* locus and a reduction in Gαs protein was demonstrated following exposure to tamoxifen (Figure 2B).

**Figure 2.**
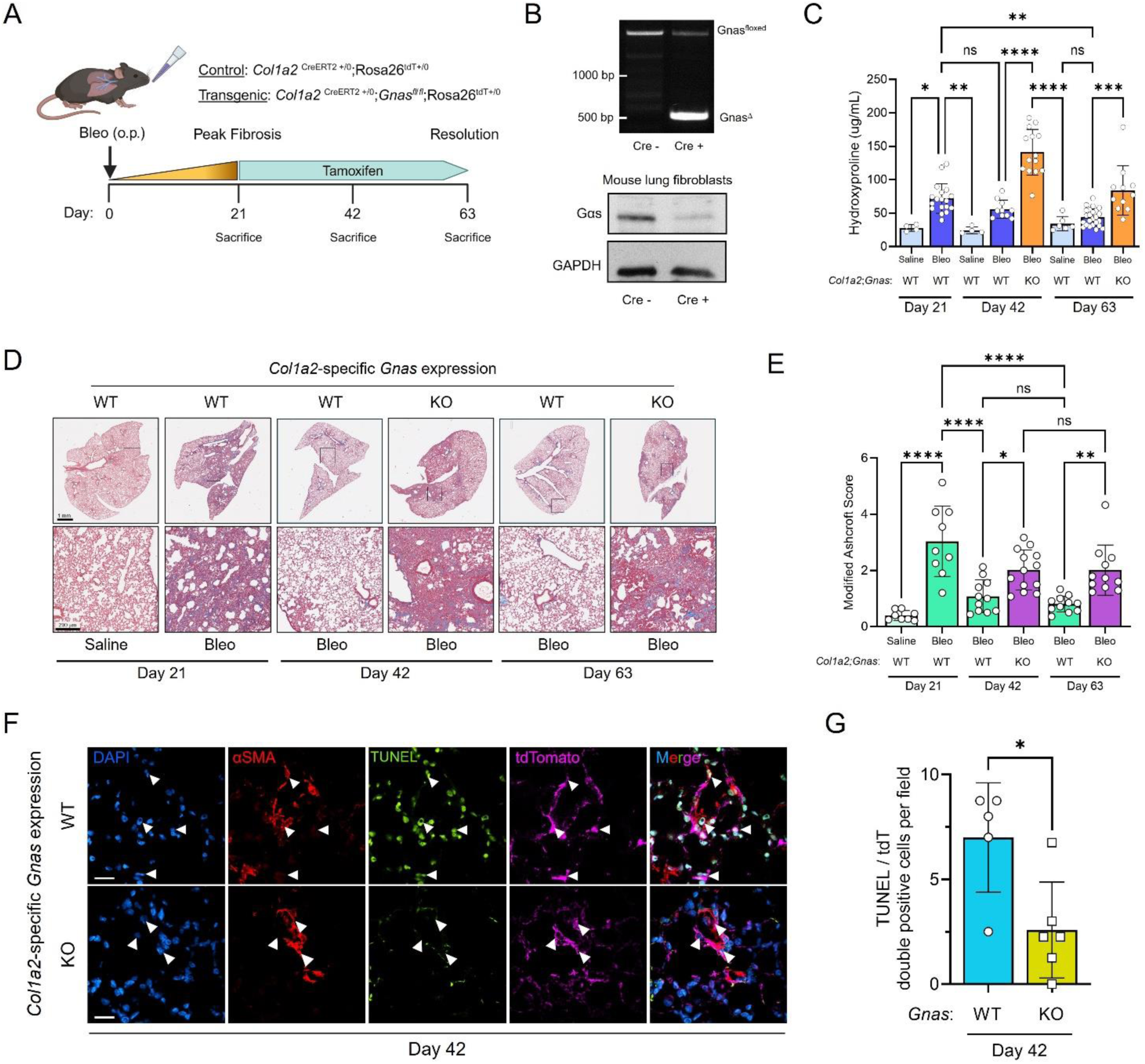
Endogenous cAMP production within lung fibroblasts following established fibrosis promotes their apoptotic clearance and is necessary for spontaneous fibrosis resolution. **(A)** Schematic illustrating the resolution protocol in WT and *Gnas* KO mice. (B) PCR of the *Gnas* locus in Cre^−^ and Cre^+^ mouse tails following tamoxifen administration (top), and subsequent Gαs protein by Western blot in Cre^−^ or Cre^+^ cultured lung fibroblasts (bottom). (**C**) Hydroxyproline content quantified from the left and right upper/middle lobe lung homogenates in saline- and bleo-treated WT lungs at day 21 and in bleo-treated lungs in WT and KO mice at days 42 and 63. (**D**) Representative images of Masson’s trichrome staining of the right lower lobe in saline- and bleo-treated WT Cre^−^ mice on day 21 and bleo-treated Cre^−^ or Cre^+^ mice on days 42 and 63. Scale bars: 1 mm (top row) and 200 μm (bottom row). (**E**) Modified Ashcroft scores of saline- and bleo-treated WT lungs at day 21 and in bleo-treated lungs in WT and KO mice at days 42 and 63. (**F**) Representative TUNEL staining of day 42 WT and KO lung sections. Scale bars: 10 μm. (**G**) Quantification of TUNEL/tdT “double positive” cells from sections in F. Each point represents the average number of TUNEL/tdT double positive cells averaged over four fields in individual mice, each of which is denoted by a separate symbol. \**P <* 0.05, \*\**P <* 0.01, \*\*\**P <* 0.001, and \*\*\*\**P <* 0.0001 were determined by 1-way ANOVA (**C** and **E**) or by 2-tailed *t* test (**G**).

As expected, control mice exhibited marked fibrotic changes at day 21 followed by evidence of spontaneous fibrosis resolution indicated by both a steady reduction in collagen content as evidenced by hydroxyproline content (Figure 2C), trichrome staining (Figure 2D), and modified Ashcroft scoring (Figure 2E) as well as histological signs of alveolar repair over the interval day 21-day 63. Crucially, *Col1a2*;*Gnas* KO mouse lungs failed to undergo spontaneous resolution, demonstrating persistent collagen deposition and histological evidence of fibrosis through day 63 (Figure 2C-E). Failed resolution in *Col1a2*;*Gnas* KO mice also correlated with persistently higher expression of α-SMA as well as the canonical pro-fibrotic fibroblast marker Cthrc1 protein through day 63 (Supplemental Figure 2C). As apoptotic clearance of pathologic fibroblasts is known to be an important feature of fibrosis resolution, we stained day 42 lung sections for TUNEL to detect cells undergoing apoptosis using tdTomato to identify fibroblasts (Figure 2F). A significantly higher number of tdTomato and TUNEL double positive cells were found in resolving WT lungs compared to persistently fibrotic *Col1a2*;*Gnas* KOs (Figure 2G).

### Longitudinal scRNA-seq of spontaneous resolving WT and persistently fibrotic *Col1a2;Gnas* KO mouse lungs

To further explore the specific role and impact of *Col1a2*-specific cAMP signaling within lung fibroblasts and its effect on other lung cell types during resolution time points, we performed scRNA-seq of control and *Col1a2*;*Gnas* KO mouse lungs within our resolution protocol (Figure 2A). In this analysis, 6 high-quality scRNA-Seq libraries encompassing on average 2151 genes and 7000 unique molecular identifiers (UMIs) per cell were used (Supplemental Figure 3A-B). A single-cell suspension was generated by collagenase digestion of whole lungs and submitted for sequencing using Illumina 10X and standard Seurat workflow. Uniform Manifold Approximation and Projection (UMAP) and Louvain clustering displayed 20 cell types (Figure 3A). Cell types were annotated using established identification markers from two mouse lung single-cell datasets (*23, 24*), revealing the presence of dendritic cells, T cells, B cells, NK cells, neutrophils, monocytes, interstitial macrophages (IM), alveolar macrophages (AM), vascular endothelial cells, lymphatic endothelial cells, aerocytes (aCap endothelial), general capillary (gCap endothelial) cells, alveolar type 2 (AT2) and type 1 (AT1) epithelial cells, club cells, ciliated cells, pericytes, smooth muscle (SM) cells, mesothelial cells, and fibroblasts (Figure 3B-C). At peak fibrosis, IMs constituted a larger proportion and AT2 cells a smaller proportion of recovered cells compared to saline-treated mice. In *Col1a2*;*Gnas* KO mice harvested on days 42 and 63, neutrophils demonstrated a marked increase in their relative proportion among total cells compared to WT controls (Figure 3C). A focused analysis of the WT controls (*Col1a2^CreERT+/0^;R26^tdT+/0^*) from this dataset – for the purpose of characterizing cellular shifts, crosstalk, and pathway enrichment during spontaneous fibrosis resolution – has been published separately (*20*).

**Figure 3.**
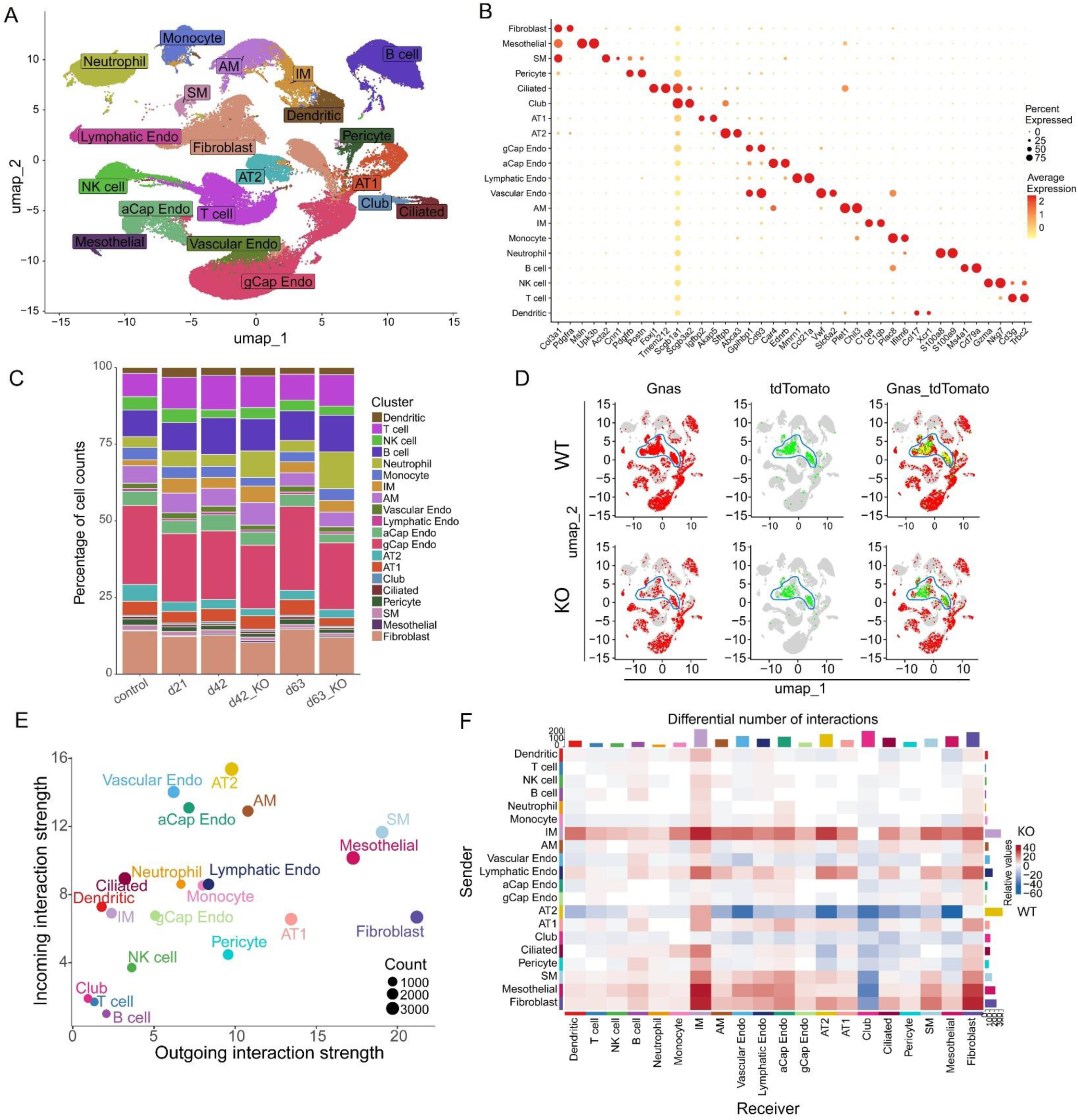
Longitudinal scRNA-seq of spontaneously resolving WT and persistently fibrotic *Col1a2;Gnas* KO mouse lungs. (**A**) Compiled UMAP plot with cell annotation of whole lung digests at each time point condition. (**B**) Scatter plot displaying top genes expressed for each annotated cell type in B. (**C**) Cluster graph displaying cell type composition at each time point. (**D**) Whole UMAP feature plots displaying *Gnas* (left), tdTomato, (middle), and overlap (right) expression in WTs (top row) and *Col1a2;Gnas* KOs (bottom row). Blue line encircles fibroblasts. (**E**) Dot plot map depicting relative incoming and outgoing interaction strength for all annotated cell types. (**F**) Heat map depicting relative number of interactions between sender cell types (y-axis) and recipient cell types (x-axis). Blue indicates an increased relative number of interactions in WT conditions while red denotes the same in KOs.

Feature plots of the global UMAP displaying relative *Gnas* and tdTomato transcript levels demonstrate the near ubiquitous expression in lung cells of the former and the expected mesenchymal restriction of the latter within WT lungs (Figure 3D). Importantly and as expected, in *Col1a2*;*Gnas* KOs, *Gnas* expression was markedly reduced in the mesenchymal compartment – correlating with tdTomato expression (Figure 3D, Supplemental Figure 3C-D).

We next employed CellChat analysis (*25*) to quantify interaction number among sender and recipient cell types as well as relative incoming and outgoing signaling strength and number of interactions for each cell population. Fibroblasts exhibited the highest overall outgoing interaction strength, while AT2, AMs and vascular/aCap endothelial cells were the recipients of the highest interaction strength (Figure 3E). Furthermore, fibroblasts and IMs were among those displaying the largest relative difference in interaction number favoring *Col1a2;Gnas* KOs (Figure 3F). Conversely, AT2 sent and club cells received more interactions in WT mice compared to KOs.

### Persistently fibrotic *Col1a2;Gnas* KO mouse lungs exhibit higher numbers of ECM-secreting *Csmd1^+^* fibroblast subtypes

To analyze the contributions of mesenchymal cells in greater detail, clusters annotated as “fibroblast,” “pericyte,” “smooth muscle (SM)”, and “mesothelial cells” from the global UMAP analysis (Figure 3A) were further subclustered (Figure 4A). Fibroblasts were identified as peribronchial, adventitial, and alveolar fibroblast subtypes, distinguished by the unique expression of *Hhip*, *Pi16*, and *Scube2*, respectively (*26*) (Figure 4B). The proportion of any single fibroblast population failed to change substantially in KO conditions compared to WT (Figure 4C).

**Figure 4.**
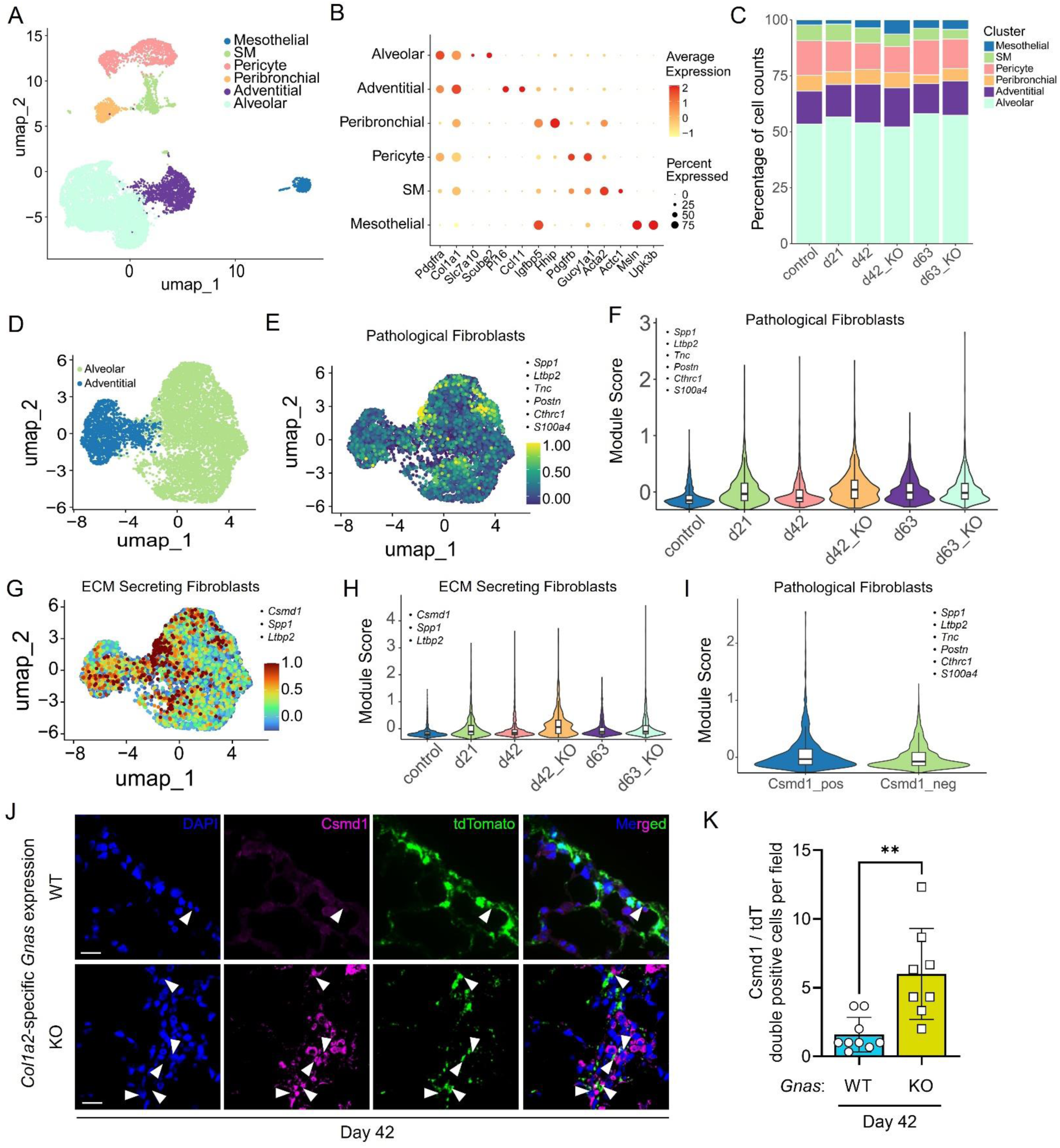
Persistently fibrotic *Col1a2;Gnas* KO mouse lungs exhibit higher numbers of ECM-secreting *Csmd1^+^* fibroblast subtypes. (**A**) UMAP of mesenchymal cell subclusters. (**B**) Dot plot of representative markers for each mesenchymal subset. (**C**) Cluster graph displaying longitudinal cell type composition of mesenchymal subtypes. (**D**) UMAP of alveolar and adventitial fibroblast subclusters. (**E**) Feature plot displaying individual cell module scores for pathological fibroblast-related genes: *Spp1*, *Ltbp2*, *Tnc*, *Postn*, *Cthrc1*, and *S100a4.* (**F**) Violin plot of module scores assigned to pathological fibroblast-related genes in E over each time point and genotype. (**G**) Feature plot displaying individual cell module scores for ECM secreting fibroblast-related genes: *Csmd1*, *Spp1*, *Ltbp2*. (**H**) Violin plot of module scores assigned to ECM secreting fibroblast-related genes in G. (**I**) Relative module score assigned to pathological fibroblast-related genes in E among *Csmd1*+ and *Csmd1*-fibroblasts. (**J**) Representative immunofluorescence microscopy images of Csmd1+ fibroblasts (determined by co-staining with tdTomato) at day 42 in WT and KO mice. White arrowheads identify Csmd1/tdT double positive cells. Scale bars: 10 μm. (**K**) Quantification of Csmd1 and tdTomato “double positive” cells in WT versus KO at day 42. \*\**P <* 0.01 was determined by 2-tailed t-test.

We next further subclustered fibroblasts into alveolar and adventitial subsets (Figure 4D) as each are located in the lung parenchyma, express tdTomato (Supplemental Figure 3D), and are known to harbor pathological fibroblast populations (*26*). Within these subsets we identified cells expressing genes associated with a pathological fibroblast phenotype (*27*) by gene module score reflecting the combined expression of *Spp1*, *Ltbp2*, *Tnc*, *Postn*, *Cthrc1*, and *S100a4* (Figure 4E). Module scores identifying pathological fibroblasts were higher in bleo-treated mice compared to saline and notably in *Col1a2;Gnas* KO mice at day 42 compared to WT controls (Figure 4F) consistent with their presumed failure to undergo apoptosis and therefore to accumulate.

Recently, a novel ECM-secreting fibroblast subpopulation identified by *Csmd1* expression was described (*28*). In this study, Guo *et al* found that *Csmd1^+^* fibroblasts expressed the most *Col1a1* and were temporally enriched during the fibrotic phase of bleo-induced injury followed by a decrease in their abundance during early resolution. Consistent with this finding, separate analysis of our dataset focused on WT mice undergoing spontaneous resolution indeed suggested an increase in ECM-secreting *Csmd1*^+^ fibroblasts at peak fibrosis – identified by a gene module score reflecting the combined expression of *Csmd1*, *Spp1*, and *Ltbp2* – followed by their decline throughout resolution (*20*). In the present analysis, *Col1a2*;*Gnas* KO mouse lung parenchymal fibroblasts exhibited higher module scores for *Csmd1*^+^ ECM-secreting fibroblasts at day 42 compared to WT controls (Figure 4G-H, Supplemental Figure 4A). Interestingly, pathological fibroblast gene signatures (Figure 4E-F) were detected in both *Csmd1*^+^ and *Cmsd1*^-^ fibroblasts, suggesting these two populations are non-nested and partially overlapping (Figure 4I). Importantly, immunofluorescence staining for Csmd1/tdT double positive cells confirmed an increased presence of this ECM-secreting fibroblast population in day 42 *Col1a2*;*Gnas* KO lung sections compared to WT controls (Figure 4J-K).

We next regrouped alveolar and adventitial fibroblasts into *Csmd1*^+^ and *Csmd1^-^* subclusters (Supplemental Figure 4B-C). Though relatively rare, the pathologic fibroblast marker transcript *Cthrc1* was detected in parenchymal fibroblasts, overlapping with *Csmd1*^+^ fibroblasts clusters (Supplemental Figure 4D). As with alveolar fibroblasts, the peribronchial cluster displayed increased *Csmd1*^+^ fibroblasts (Supplemental Figure 4E). Similar to the Guo *et al.* report describing the *Csmd1*+ population (*28*), we found that peribronchial fibroblasts lacked *Pdgfra* co-expression (Supplemental Figure 4F). ECM-secreting fibroblasts predicted by gene module score in the peribronchial compartment did not demonstrate the accumulation observed in alveolar and adventitial fibroblasts (Supplemental Figure 4G-J). Lastly, we confirmed that *Csmd1*+ fibroblasts demonstrated increased collagen gene expression (Supplemental Figure 5). Taken together, persistent fibrosis within *Col1a2*;*Gnas* KO lungs was associated with higher enrichment of pathological and *Csmd1*+ ECM-secreting fibroblast clusters present within the alveolar/adventitial fibroblast populations.

### Fibrosis-associated macrophages persist in fibrotic *Col1a2;Gnas* KO mouse lungs

Lung macrophage populations are known to be important in fibrotic responses (*29*). Indeed, their depletion with clodronate during the fibrotic phase of bleomycin-induced lung injury has been shown to abrogate fibrosis (*30*), attesting to their pro-fibrotic potential. Furthermore, a number of studies have demonstrated the accumulation of monocyte-derived macrophages following injury that adopt IM and/or AM markers and express pro-fibrotic genes (*31–33*). Though the effect of cAMP signaling within lung macrophages is well known to be inhibitory (*34*), it is not known if fibroblast-intrinsic cAMP signaling can influence these cells during resolution. To more precisely characterize changes in lung macrophage subsets in resolving versus persistent fibrosis within *Col1a2*;*Gnas* KO mice, clusters annotated as “AM” and “IM” from our global UMAP (Figure 3A) were further subclustered.

We identified four distinct populations based on established gene markers (Figure 5A-B). Tissue resident AMs (TRAMs) exhibited high expression levels of *SiglecF*, *Ear1*, *Car4*, *Cd11c* (Itgax) and Fabp5 (Figure 5B, Supplemental Figure 6A). IMs were distinguished from AMs by elevated levels of *CD11b* (Itgam), *Cx3cr1*, and *C1qa*. A proliferating TRAM (pro-TRAM) population was identified by high concomitant expression of *Stmn1* and *Mki67*. Finally, a cluster co-expressing both IM and AM markers (Figure 5B) along with several pro-fibrotic genes – *Arg1*, *Spp1*, *Trem2*, *Cd63*, *Fabp5*, and *Gpnmb* (Figure 5D) – appeared at peak fibrosis and persisted through the subsequent weeks in *Col1a2*;*Gnas* KOs (Figure 5C & E, Supplemental Figure 6B). This population, which we previously termed fibrosis-associated macrophages (FAMs) (*20*), is consistent with those described in previous studies of bleo-induced lung injury (*31, 35*). Heatmap of top expressed genes in macrophage subsets (Supplemental Figure 6A) displays genes further consistent with expression previously reported in this population. In contrast to their decline during resolution in WT mice by relative cell abundance (Figure 5C) and gene module score (Figure 5E), FAM-related genes remained markedly elevated on day 42 and with persistent elevation through day 63 in KO mice. Consistent with this finding, immunofluorescence staining confirmed the increased presence of Arg1+Cd63+ macrophages within lung sections of KO mice obtained 42 days post-bleo (Figure 5F-G).

**Figure 5.**
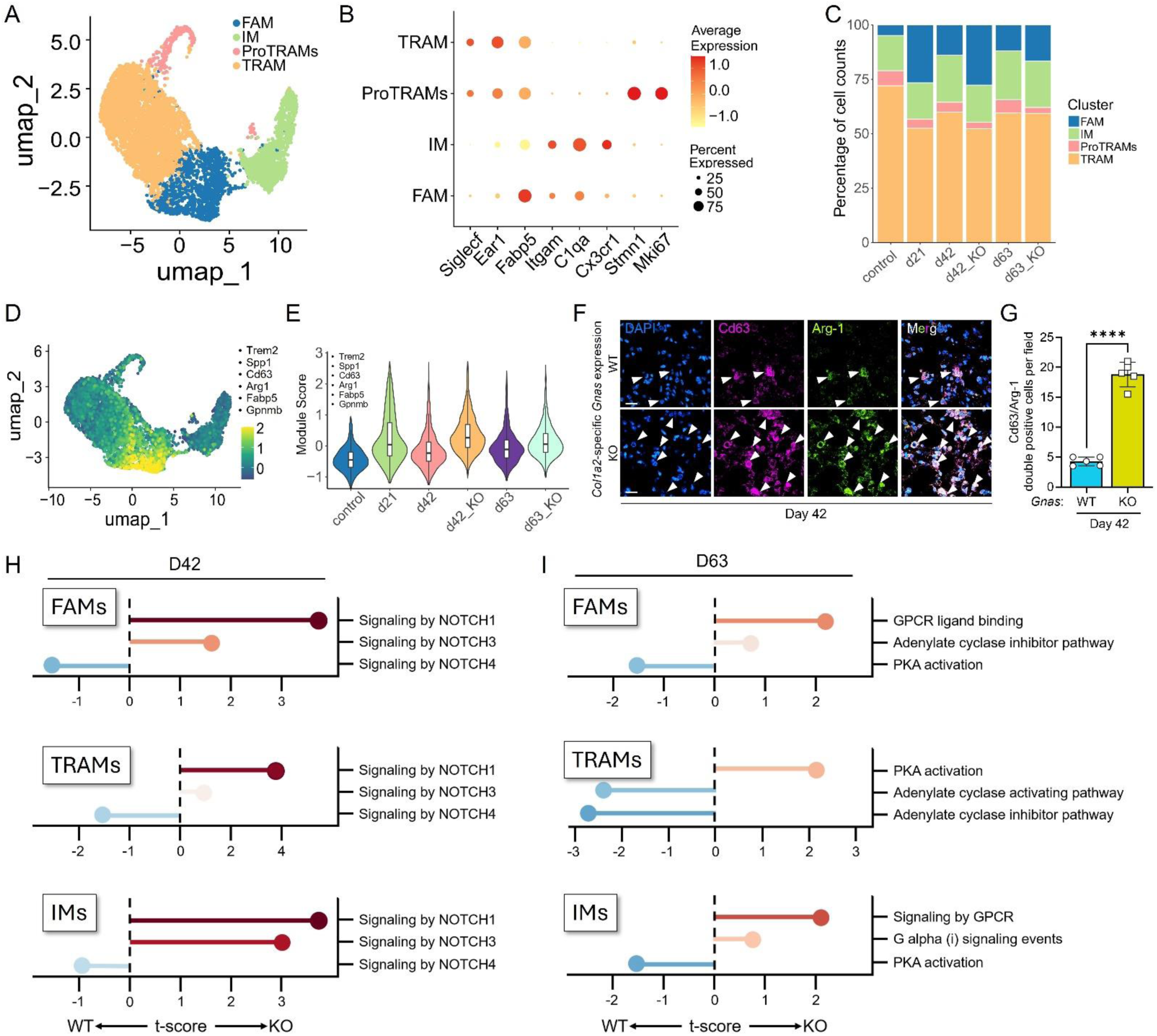
Fibrosis-associated macrophages persist in fibrotic *Col1a2;Gnas* KO mouse lungs. (**A**) UMAP of macrophage subclusters displaying tissue resident alveolar macrophages (TRAMs), interstitial macrophages (IMs), proliferating macrophages (ProTRAMs), and fibrosis-associated macrophages (FAMs). (**B**) Dot plot of representative markers for each macrophage subtype. (**C**) Cluster graph displaying longitudinal cell type composition of macrophage subtypes. (**D**) Feature plot displaying individual cell module scores for FAM-related genes:*Trem2*, *Spp1, Cd63, Arg1, Fabp5, Gpnmb*. (**E**) Violin plot of module scores assigned FAM-related signatures genes in D over each time point and genotype. (**F**) Representative immunofluorescence staining of Cd63 and Arg1 in FAMs from WT and KO lungs at day 42. Scale bars: 10 μm. (**G**) Quantification of Arg1 and Cd63 “double positive” cells in WT versus KO at day 42. (**H**) Waterfall plots depicting relative enrichment of NOTCH pathways in day 42 WT versus KO FAMs, TRAMs, and IMs. (**I**) Waterfall plots depicting relative enrichment of cAMP-related pathways and downstream signaling in day 63 FAMs, TRAMs, and IMs. \*\*\*\**P <* 0.0001 was determined by 2-tailed t-test.

Our previous longitudinal GSEA in WT mice (*20*) revealed enrichment of the inflammatory-associated Notch 1 and fibroblast-associated Notch 3 signaling pathways in FAMs and AMs at day 21 followed by reduced enrichment of each during resolution. The vascular-associated Notch 4 pathway showed a complementary pattern with its highest enrichment within each subtype evident in saline-treated lungs compared to peak fibrosis. In the current study, GSEA comparisons were performed between WT and KOs, revealing enrichment in Notch 1 and 3 signaling within FAMs, AMs, and IMs from *Col1a2;Gnas* KO lungs compared to controls at day 42 (Figure 5H). In contrast, Notch 4 was enriched in each subset from WT lungs on day 42. Thus, distinct Notch pathways within macrophage subsets demonstrated variable or even opposing patterns of enrichment in resolving WT and non-resolving Gnas KO lungs.

We previously reported macrophage subsets in WT mice that demonstrated enrichment in pathways that inhibit cAMP production at peak fibrosis with a shift towards pathways favoring cAMP signaling during spontaneous resolution (*20*). To assess whether cAMP signaling within lung fibroblasts influences this same second-messenger pathway in macrophages during resolution, we once again compared GSEA in WT to that in *Col1a2;Gnas* KO mice (Figure 5I). Both FAMs and IMs from *Col1a2;Gnas* KO lungs exhibited enrichment in cAMP inhibitory pathways at day 63 while in WT lungs these cells were enriched for PKA activation (Figure 5I). Surprisingly, an opposite pattern was demonstrated within AMs from *Col1a2;Gnas* KOs, reflecting differences among subtypes within resolving and persistently fibrotic lungs.

### Persistently fibrotic *Col1a2*;*Gnas* KO lungs display abnormal epithelial remodeling with altered repair pathway enrichment

Impaired alveolar regeneration is a hallmark of persistent/progressive pulmonary fibrosis. In our model, *Col1a2;Gnas* KO mouse lungs displayed histopathology consistent with abnormal epithelial repair in fibrotic regions (Figure 2D). To gain further insight into the distinct epithelial cell phenotypes and pathway enrichment patterns within resolved versus persistently fibrotic lungs, clusters annotated as “AT1”, “AT2”, “ciliated”, and “club” cells were subclustered from the global UMAP and reanalyzed using cell specific markers (Figure 6A-B). In addition to the aforementioned cell types, we identified a cluster of epithelial cells aligned between AT1 and AT2 cells expressing *Cldn4* and *Sfn*. This “transitional” cell population was not present in controls and constituted a small proportion of epithelial cells at day 21 that declined during resolution in WT mice (Figure 6C). In *Col1a2;Gnas* KOs, transitional cells displayed a higher frequency at day 42 but declined to levels consistent with WT lungs by day 63. The proportion of the remaining epithelial cells did not display substantial differences longitudinally (Figure 6C).

**Figure 6.**
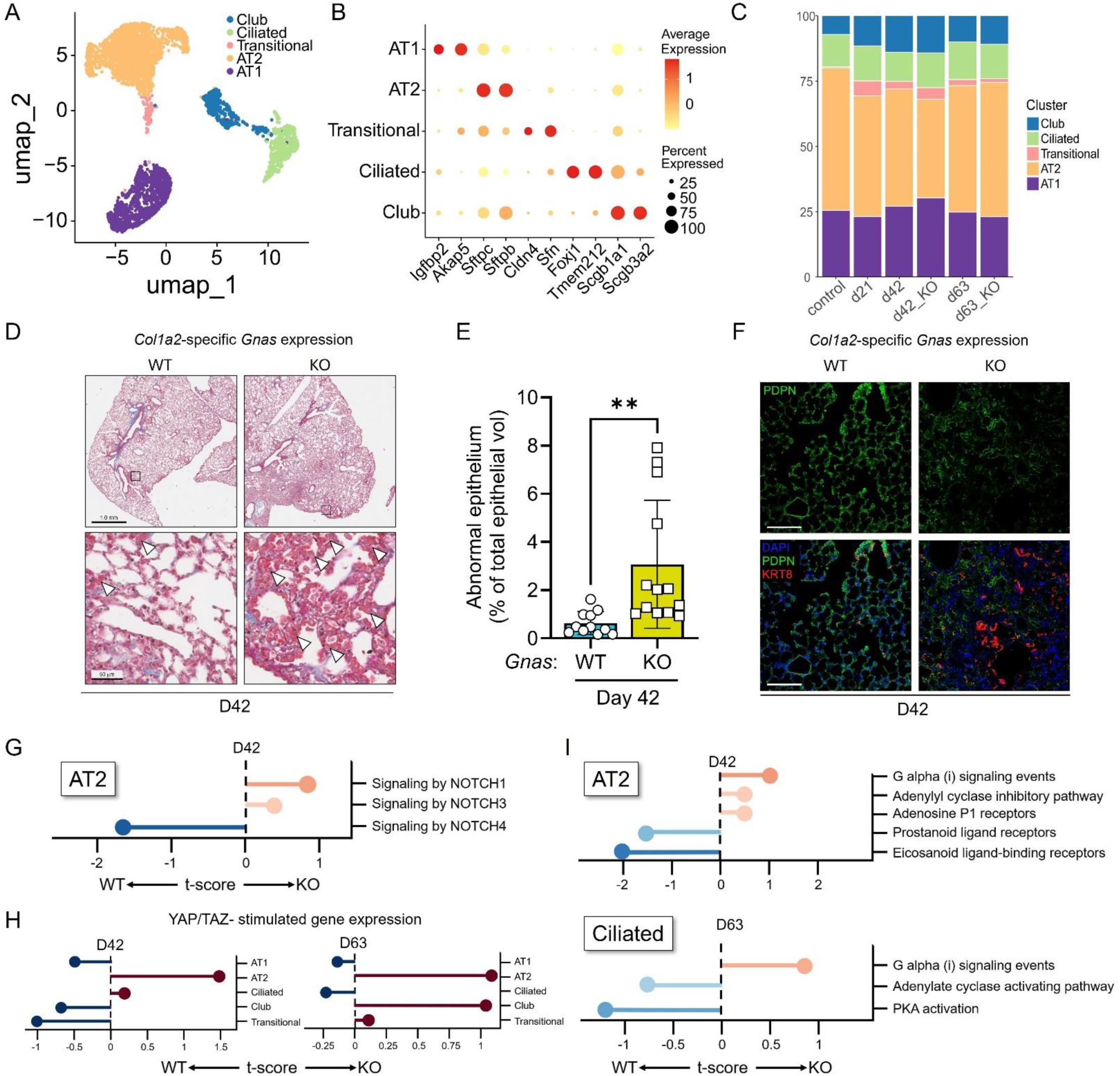
Persistently fibrotic *Col1a2*;*Gnas* KO lungs display abnormal epithelial remodeling with altered repair pathway enrichment. (**A**) UMAP of epithelial cell subclusters. (**B**) Dot plot of representative markers for each epithelial subset. (**C**) Cluster graph displaying longitudinal cell type composition of epithelial subtypes. (**D**) Representative images of Masson’s trichrome staining of the right lower lobe. Arrowheads identify regions of abnormal cuboidal alveolar epithelial cells. Scale bars: 1.0 mm (top row) and 50 μm (bottom row). (**E**) Quantification of abnormal cuboidal epithelial cells from lung sections in D. Each point represents the average percentage of abnormal epithelial among total epithelial cells quantified from lung sections obtained from different mice. (**F**) Immunofluorescence staining of type I alveolar epithelial (PDPN) and keratin 8 (Krt8) positive cells in WT and KO day 42 lung sections. Scale bars: 100 μm. (**G**) Waterfall plot depicting relative AT2 NOTCH signaling enrichment in WT versus KOs at day 42. (**H**) Waterfall plots depicting relative YAP/TAZ gene expression enrichment in day 42 and 63 WT versus KOs among epithelial cell subtypes. (**I**) Waterfall plots of depicting relative enrichment of cAMP-related pathways and downstream signaling in day 42 AT2 cells and day 63 ciliated airway cells in WT versus KOs. \*\**P <* 0.01 was determined by 2-tailed t-test.

Despite the relative paucity of transitional cells during peak fibrosis and resolution compared to their elevated proportions during the inflammatory phase of lung injury and early fibrogenesis (*36*) – which we further confirmed by CLDN4 staining (Supplemental Figure 7) – *Col1a2;Gnas* KO lungs exhibited scattered regions of abnormal epithelial remodeling in which alveolar spaces were lined with cuboidal epithelial cells (Figure 6D-E). We next stained for Krt8, a broad marker of intermediate cell populations, and found disorganized clusters of parenchymal Krt8+ cells in regions devoid of the AT1 marker PNPD (Figure 6F). The airway-like structures formed by Krt8+ cells within the parenchymal regions of *Col1a2;Gnas* KO lungs are consistent with bronchiolization or AT2 cell hyperplasia – pathologic features observed in non-resolving models of fibrosis (*37*) and human pulmonary fibrosis (*38*).

Several pathways related to alveolar regeneration and development have been shown to be dysregulated in persistent fibrosis. Among them, Notch and Hippo-Yap/Taz signaling have each been associated with bronchiolization as well as AT2 hyperplasia (*39–42*). We therefore assessed Notch and Hippo pathways by GSEA within AT2 and bronchiolar epithelial cells. Similar to the pattern observed in lung macrophages, AT2 and ciliated cells demonstrated enrichment in Notch 1 and 3 pathways in *Col1a2;Gnas* KOs while Notch 4 tended to be enriched in the control mice (Figure 6G). Gene expression related to the Hippo pathway transcriptional co-activators Yap and Taz was found to be enriched within AT2 cells in *Col1a2;Gnas* KOs on days 42 and 63 as well as in club cells from KOs on day 63 (Figure 6H). Complementing these findings, cAMP-related signaling – which can inhibit Yap/Taz signaling (*43*) – was relatively suppressed in AT2 and Club cells within KOs and enriched in WT controls on days 42 and 63 with a similar pattern in ciliated cells on day 63 (Figure 6I).

### Lung fibroblast cAMP signaling influences pro-resolution crosstalk among immune, epithelial, and mesenchymal cells during spontaneous resolution

We performed CellChat analysis to map signaling interactions among selected mesenchymal (alveolar and adventitial fibroblasts), immune (FAMs, TRAMs, IMs), and epithelial (AT1 and AT2) cell subtypes during resolution in both WT and *Col1a2;Gnas* KO mice. Total interaction number and strength among cell types analyzed was highest in WT bleo-treated lungs at day 21 with a gradual decline through day 42 and 63 in WT and KOs (Supplemental Figure 8A). Interaction strength was greater in day 63 KO compared to day 63 WT lungs, with small relative differences in outgoing and incoming signal strength of each cell type contributing to this difference (Supplemental Figure 8B). Further delineation of cell-cell signaling was determined by quantifying the number of interactions between each cell type. On day 21, FAMs served as both the source and recipient cell for substantially more interactions in bleo-treated lungs than in saline controls, owing to their massive expansion/recruitment during fibrosis (Figure 7A). In day 42 bleo-treated KOs, FAMs received more interactions from parenchymal fibroblasts and AT2 cells and sent more signals to alveolar fibroblasts than in WT lungs (Figure 7B). TRAMs displayed the opposite recipient pattern at day 42, receiving more interactions from all cell types in WT lungs. Among epithelial cells, AT2 cells from KO lungs sent more signals to FAMs, IMs, AT1 cells, and adventitial fibroblasts at day 42 compared to day 63 (Figure 7C). As a recipient cell, AT2s received more signals at day 63 from AT1 and alveolar fibroblasts in KO lungs. Finally, AT1 cells in KO lungs were the source of a greater number of outgoing interactions by day 63.

**Figure 7.**
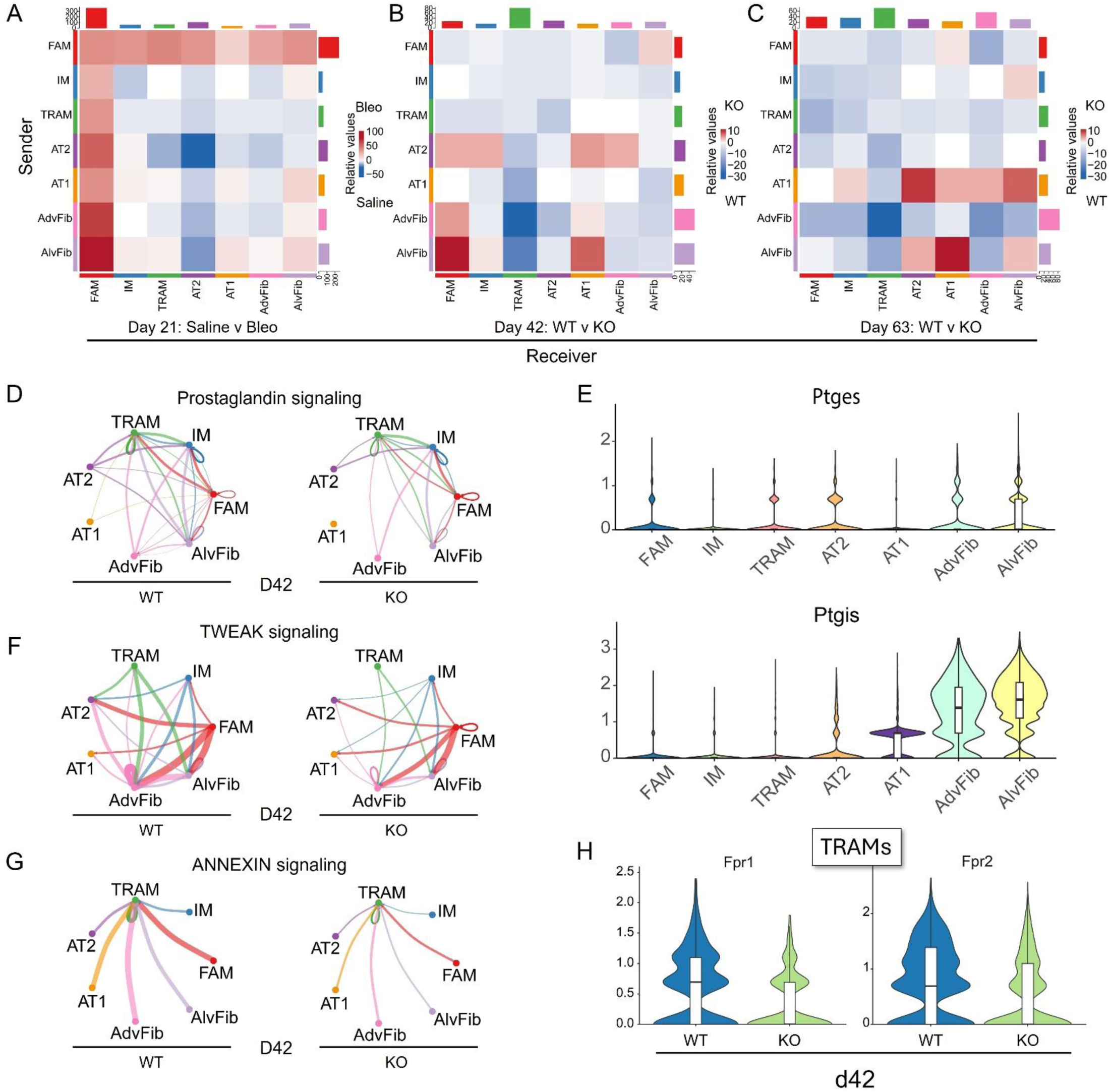
Lung fibroblast cAMP signaling influences pro-resolution crosstalk among immune, epithelial, and mesenchymal cells during spontaneous resolution. **(A – C)** Heatmaps depicting relative number of interactions between sender cell types (y-axis) and recipient cell types (x-axis). Blue indicates an increased relative number of interactions in saline (A) or WT conditions (B and C) while red denotes the same in bleo (A) or KO (B and C). (D, F, G) Circos plots depicting cellular crosstalk among the indicated pathways in WT and KO lungs at day 42. (D) Prostaglandin signaling network and relative expression of PGE_2_ synthase (*Ptges*) and PGI_2_ synthase (*Ptgis*) expression by cell type (E). (F) TWEAK signaling network in day 42 WT and KOs. (G) Annexin signaling network and relative annexin receptor expression in day 42 TRAMs from WT and KO lungs (H).

As cellular crosstalk is likely to be crucial for normal lung repair and fibrosis resolution, we measured the relative global information flow among the aforementioned cell types in control and *Col1a2;Gnas* KO lungs at day 42 and 63 (Supplemental Figure 8C). Several pro-fibrotic pathways involved in inflammation, fibrosis, and tissue repair demonstrated differences between conditions. For example, SPP1 and IL1 were enriched in KOs at each time point while KOs displayed more collagens, FN1, and BMP pathway signaling on day 63. Specifically, SPP1 exhibited more TRAM- and FAM-mediated cellular crosstalk in KOs, particularly at day 42 (Supplemental Figure 9A). However, pathways such as TGFβ and MIF did not display substantial differences between WT and KOs and indeed displayed sustained increases in cellular crosstalk in resolved lungs (Supplemental Figure 9B-C). As alterations in pro-resolution signaling may account for differences between persistent and resolved fibrosis in the setting of sustained pro-fibrotic factor exposure, we next assessed cell-cell communication involving putative anti-fibrotic pathways.

Several pathways with anti-fibrotic properties such as prostaglandin, annexin, and TWEAK/Fn14 were found to be enriched during resolution in WT lungs compared to KOs by information flow (Supplemental Figure 8C). Specifically, prostaglandin signaling at day 42 in KO mice demonstrated reduced crosstalk between epithelial and immune/fibroblasts as well as fibroblasts with immune and epithelial cells (Figure 7D). The prostaglandins encompass a subgroup of eicosanoid lipid mediators that signal via GPCRs (*44*). Among them, PGE_2_ and PGI_2_ notably signal via cAMP and can exhibit anti-fibrotic/pro-resolution properties within lung fibroblasts (*45, 46*), epithelial (*47*), and immune cells (*48, 49*). Gene expression analysis of the prostaglandin pathway using CellChat revealed GPCR receptor substantial expression of the terminal PGI_2_ biosynthetic enzyme PGI_2_ synthase (Ptgis) in alveolar and adventitial fibroblasts, dwarfing its expression in other cell types and the expression levels of the terminal PGE_2_ biosynthetic enzyme Ptges (Figure 7E, Supplemental Figure 10A). Notably, transcript expression of the PGI_2_ receptor *Ptgir* was found to be reduced within FAMs from *Col1a2*;*Gnas* KOs compared to WT mouse lungs at day 63 (Supplemental Figure 10B).

Another soluble factor influencing fibrotic outcomes through crosstalk is the cytokine TWEAK, which has been shown to exhibit anti-fibrotic properties in pulmonary fibrosis through its signaling in fibroblasts (*50*). TWEAK showed complex crosstalk, originating from immune and mesenchymal cells and signaling to fibroblasts and epithelial cells, with a substantial reduction in KOs at day 42 (Figure 7F).Transcript levels of the TWEAK receptor (*Fn14*) were previously predicted to increase in parenchymal fibroblasts during spontaneous resolution (*20*) and displayed a modest reduction in expression within 63 KO lungs compared to WT controls in the current study (Supplemental Figure 10C). TRAMs were found to be the sole recipient cell for annexin signaling with source contributions from each other cell type (Figure 7G). Annexin crosstalk was predicted to be universally reduced in day 42 KOs compared to WT controls. Furthermore, expression of Fpr1 and Fpr2 receptors (which meditate annexin signaling via Anxa1 binding) were reduced in TRAMs in KO lungs (Figure 7H).

### cAMP signaling within dermal fibroblasts is necessary for spontaneous resolution of skin fibrosis

Anti-fibrotic effects related to cAMP signaling have been demonstrated to extend beyond the lungs (*51–54*). For example, dermal myofibroblasts contributing to skin fibrosis – a highly morbid manifestation of systemic sclerosis/scleroderma – can be dedifferentiated *in vitro* by increasing intracellular cAMP (*55*). To determine whether fibroblast cAMP signaling is necessary to resolve fibrosis in the skin as it is in the lung, we employed a well-established mouse model of skin fibrosis, utilizing subcutaneous administration of bleo; this model mimics certain key features of scleroderma (Figure 8A), yet has been established to exhibit spontaneous resolution (*56*). Bleo (10 mg/kg) or an equal volume of saline was administered by subcutaneous injection to WT and *Col1a2^CreERT2+/0^;Gnas^fl/fl^* female mice daily for 7 consecutive days. Mice were sacrificed and skin samples were harvested on day 24 or 63 to capture peak fibrosis and resolution, respectively. Mice from the day 63 cohort were provided with a tamoxifen chow diet starting on day 24. Timepoints for harvest were selected based on published literature (*57*) and further refined by pilot experiment readouts for peak fibrosis and resolution (data not shown). Fibroblasts were cultured from skin obtained from WT and *Col1a2;Gnas* KO mice to demonstrate that Cre-mediated recombination of *Gnas* (confirmed by PCR of tail snips) reduced Gαs protein expression by Western blot (Figure 8B).

**Figure 8.**
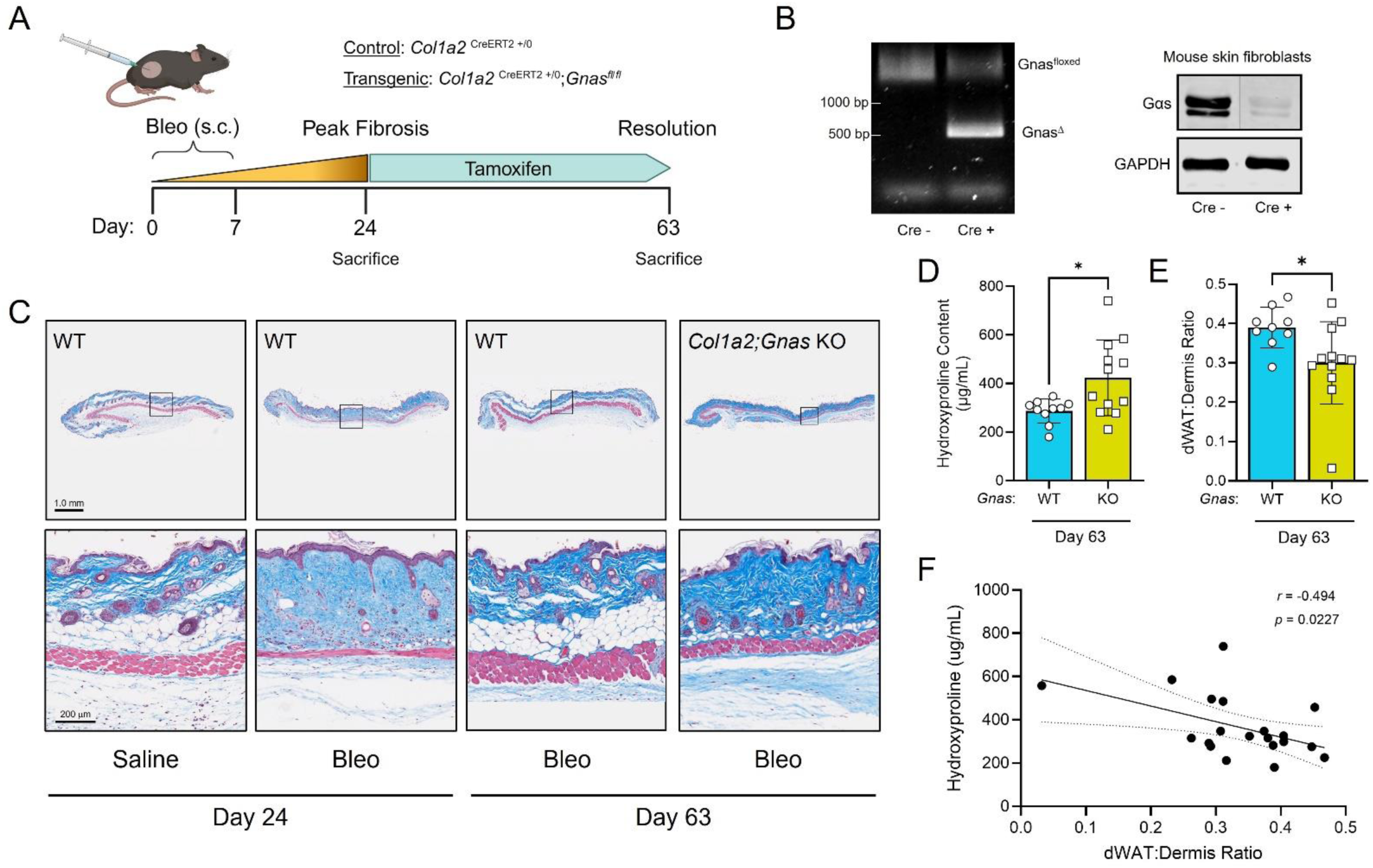
cAMP signaling within dermal fibroblasts is necessary for spontaneous resolution of skin fibrosis. (**A**) Schematic of subcutaneous bleo model and spontaneous skin resolution protocol. (**B**) PCR of the *Gnas* locus in Cre^−^ and Cre^+^ mouse tails following tamoxifen administration (left), and subsequent Gαs protein by Western blot in Cre^−^ or Cre^+^ cultured skin fibroblasts (right). (**C**) Representative images of Masson’s trichrome staining of the skin in saline- and bleomycin-treated WT Cre^−^ mice on day 24 and bleo-treated Cre^−^ or bleo-treated Cre^+^ (*Col1a2*;*Gnas* KO) mice on day 63. Scale bars: 1.0 mm (top row) and 200 μm (bottom row). Hydroxyproline content (**D**) and dWAT:dermis ratio (**E**) quantified from day 63 WT and *Gnas* KO skin sections. (**F**) Linear regression analysis of dWAT:dermis ratio measured in WT and KO d63 lung sections compared to hydroxyproline content. The correlation coefficient “r” was determined using a Pearson correlation test. \**P <* 0.05 was determined by 2-tailed t-test.

Skin histopathology from bleo-treated mice displayed a thickened dermis and increased collagen by trichrome staining on day 24 (Figure 8C). Importantly, a substantial reduction in dermal white adipose tissue (dWAT) thickness was also observed in bleo-treated mice, consistent with the pathology of skin fibrosis in scleroderma (*58, 59*). By day 63, bleo-treated WT mice exhibited a thinner dermis compared to day 24 with a reappearance of the dWAT layer characteristic of resolution. In contrast, *Col1a2;Gnas* KO skin displayed higher average dermal thickness and hydroxyproline content compared to WT with a thinner dWAT and lower dWAT:dermal ratio (Figure 8C-E). A lower dWAT:dermis ratio was found to correlate with greater hydroxyproline content (Figure 8F).

## DISCUSSION

Persistent/progressive fibrosis develops following tissue injury when pathways responsible for halting the fibrotic response and reestablishing cellular homeostasis are disabled. Animal models characterized by spontaneous resolution of fibrosis can be leveraged to identify endogenous pathways inactivated in states of fibrosis whose restoration may have therapeutic potential. Within the lung, there is now ample evidence in animal models to suggest that failed dedifferentiation/clearance and thus accumulation of apoptosis-resistant pathologic fibroblasts/MFs represents a major determinant of persistent fibrosis after injury (*6, 8, 12, 22*). Indeed, in addition to their essential role in parenchymal scarring, persistence of these cells is associated with macrophage activation and dysregulated epithelial repair – cardinal features of IPF (*22, 60*). A rigorous body of research spanning several decades has demonstrated both the robust and pleiotropic inhibitory properties of cAMP on lung fibroblast activation (*61–64*), as well as numerous defects within the cAMP signaling axis in fibrotic lung diseases (*65–67*). In addition to inhibiting multiple pro-fibrotic driver pathways, cAMP also increases the expression of a variety of anti-fibrotic brakes (*19*) within lung fibroblasts – including the transcription factor KLF4 and the protein phosphatase MKP1 – which have themselves each been shown to promote MF dedifferentiation and restore their apoptosis sensitivity (*22, 68*). However, the *in vivo* requirement for resolution of endogenous cAMP signaling specifically within lung fibroblasts has not previously been investigated. In the present study, we addressed this knowledge gap by employing fibroblast-specific Gαs loss-of-function to show that endogenous fibroblast cAMP production following the establishment of lung fibrosis (i) promotes fibroblast deactivation/dedifferentiation and apoptosis, (ii) is necessary for fibrosis resolution, and (iii) orchestrates alveolar regeneration and immune cell homeostasis through fibroblast-epithelial and fibroblast-macrophage crosstalk. Furthermore, we confirmed that the pro-resolution effects of intracellular cAMP within fibroblasts likewise extend beyond the lung to skin fibrosis. To our knowledge, this study is the first to abolish the capacity of receptor-mediated cAMP generation in fibroblasts from any organ or in resident lung cell types of any kind in order to define its role during fibrosis resolution. Our findings establish the importance of fibroblast cAMP signaling during fibrosis resolution and provide mechanistic insight into the potential benefit of new (the phosphodiesterase 4B inhibitor nerandomilast) and imminent (the PGI_2_ analog treprostinil) cAMP-based therapies for IPF.

Our choice to delete *Gnas* in fibroblasts only after fibrosis had already been established allowed us to interrogate the importance of cAMP signaling within these cells specifically and exclusively during the resolution phase. Fibroblasts express a multitude of Gαs-coupled GPCRs (*69*), and the specific endogenous ligand(s) that serve to stimulate cAMP production during fibrosis resolution, their cellular source(s), and the precise timing of this signaling remain to be determined for both the lungs and the skin. Recent reports of the efficacy of treprostinil when administered to patients with IPF (*70, 71*) support the possibility that its endogenous congeners PGI_2_ and/or PGE_2_ may contribute to these anti-fibrotic effects in the lung. In our model, parenchymal lung fibroblasts were the chief source of PGI_2_ synthase transcript expression (Figure 7E), highlighting their capacity to mediate autocrine and paracrine cAMP signaling. Fibroblast-specific *Gnas* KO lungs demonstrated a distinct increase in collagen content (Figure 2C), reduced fibroblast/MF apoptosis (Figure 2F-G), and reduced pro-resolution pathway enrichment (Figure 7D-H) by day 42 – suggesting that the crucial window for cAMP production in these cells occurs within the first 2-3 weeks following peak fibrosis. Indeed, previous studies have demonstrated global transcriptomic changes in fibroblasts from spontaneously resolving lungs by day 30 post-bleo (*72*).

Persistent fibrosis in our *Col1a2*;*Gnas* KO mouse model was associated with enrichment in pathological fibroblast signatures (Figure 4E-F) as well as elevated numbers of ECM-secreting *Csmd1^+^* fibroblasts (Figure 4G-K). This cell subtype was identified and characterized by Guo *et al*. using a unique approach in which histological signatures of matrix architecture guided subsequent single-cell characterization (*28*). *Csmd1^+^*fibroblasts were found to be highest during peak fibrosis and to gradually decline during spontaneous resolution, in line with our recent (*20*) and current (Figure 4) findings. Furthermore, *Csmd1* expression within ECM-secreting fibroblasts correlated with co-expression of *Pdgfra* which we similarly found to be absent in peribronchial fibroblasts but present in alveolar and adventitial fibroblasts (Supplemental Figure 4E-F). It is notable that the small number of alveolar fibroblasts expressing *Cthrc1* overlapped with the UMAP region of highest *Csmd1* expression in this cluster (Supplemental Figure 4D). Guo *et al.* additionally described enrichment of *Cthrc1* within *Csmd1^+^* fibroblasts and demonstrated a strong alignment of Cthrc1 protein expression within Csmd1^+^ fibroblasts in IPF lungs (*28*). The persistence of ECM-secreting *Csmd1^+^* parenchymal fibroblasts in *Col1a2*;*Gnas* KO mouse lungs suggests that cAMP signaling is necessary to clear this pathologic fibroblast cell type.

An unexpected finding was that in addition to its direct effects within mesenchymal cells, abrogation of cAMP signaling in fibroblasts also impacted macrophage as well as epithelial cell phenotypes and pathway enrichment. With respect to macrophages, this was most evident in the persistence of the FAM population at day 42 – a population likely derived from recruited monocytes based on similar surface markers, gene expression, and UMAP alignment to those found in other studies (*31, 35*). FAMs exhibited the highest strength of outgoing interactions among immune cells and the greatest differential number of interactions in *Col1a2*;*Gnas* KOs, suggesting that they serve an important role in cellular crosstalk in states of persistent fibrosis. Indeed, the high interaction number likewise found in IMs from *Col1a2*;*Gnas* KOs (Figure 3F) likely reflects the contribution of FAMs before subclustering was performed, given that FAMs share similar genes used to annotate IMs. Enrichment in Notch signaling in FAMs, along with TRAMs and IMs, in *Col1a2*;*Gnas* KO lungs is consistent with a pro-fibrotic phenotype (*31, 73*), as is the parallel reduced enrichment of the antifibrotic annexin pathway (Figure 7). The effect of cAMP within lung macrophages *in vivo* is expected to be pro-resolving given its anti-inflammatory and inhibitory effects (*34*). Remarkably, we found that abolishing cAMP signaling within fibroblasts unexpectedly also diminished its signaling within FAMs and IMs (Figure 5I). In contrast, TRAMs displayed evidence of enriched cAMP signaling in KOs and a mix of activating and inhibitory inputs on this pathway in controls. Additional studies will be required to further characterize the cell-specificity and timing of cAMP activation among immune cells and their crosstalk with mesenchymal cells in persistent and resolving fibrosis.

Persistent fibrosis within *Col1a2*;*Gnas* KO lungs was also characterized by significant increases in abnormal epithelial regeneration characterized by AT1-devoid regions lined with cuboidal alveolar epithelium, as compared to the reconstitution of normal-appearing alveolar structure by AT1 cells in controls (Figure 6D-E). The Krt8^+^/Cldn4^-^ cells accumulating in persistently fibrotic lungs are likely to represent bronchiolization and/or AT2 cell hyperplasia. Both forms of dysregulated epithelial repair are considered maladaptive as these regions are poorly suited for gas exchange. Bronchiolization represents ectopic airway (ciliated or club cells) differentiation or migration and is thought to contribute to honeycomb cyst formation in IPF, whereas AT2 hyperplasia is associated with ongoing tissue remodeling and fibrosis (*60*). Recent work has suggested that sustained activity of the Hippo-Yap/Taz developmental pathway can promote aberrant AT2 cell accumulation (*40*) as well as drive bronchiolization (*42*). Our findings that AT2 and bronchiolar epithelial cells from *Col1a2*;*Gnas* KO lungs were enriched in both Yap/Taz and Notch signaling (Figure 6G-H) suggest that lung fibroblasts, via these pathways, influence epithelial cell fate. The specific mechanism by which fibroblasts exert this influence is unclear, but it is notable and equally remarkable that, as for macrophages, cAMP signaling was also predicted to be abrogated in epithelial populations such as AT2 and ciliated cells from *Col1a2*;*Gnas* KOs (Figure 6I).

This is the first study of which we are aware to demonstrate that endogenous cAMP signaling within fibroblasts is crucial for the spontaneous fibrosis resolution observed following limited tissue injury to the lungs and skin. Our findings raise a number of questions related to the actions of cAMP in the context of tissue fibrosis. These include whether cAMP represents a ubiquitous resolver of fibrosis following injury to other organs (i.e. liver, kidney, heart, etc.) as well as its specific *in vivo* role in non-mesenchymal cell types. These questions can be approached by using *Gnas^fl/fl^* mice strategically coupled to cell-specific Cre recombinases within spontaneously resolving fibrosis models appropriate to the organ of interest. Furthermore, putative anti-fibrotic therapies that act downstream of cAMP can be tested for their efficacy to resolve persistent fibrosis in *Col1a2;Gnas* KO mice. Beyond animal models, validation of the role of cellular cAMP machinery within healthy and fibrotic human tissue will be necessary. Lastly, the mechanism by which fibroblast-specific cAMP abrogation might lead to the diminution of cAMP signaling in FAMs, IMs, AT2, and bronchiolar cells, as suggested by our scRNA-seq analysis, remains an open question. Fibroblasts may serve as a major source of cAMP-producing ligands, including PGI_2_ and PGE_2_, for epithelial and immune cells whose synthesis and secretion are themselves cAMP dependent (*74*). Alternatively, effects on cAMP signaling in neighboring cell types could be mediated either by direct cAMP efflux and intercellular transfer through connexins (*75*), or its efflux via ABCC family transporters with subsequent extracellular conversion to adenosine acting via Gαs- and Gαi-coupled receptors (*76*). Answering these questions will provide a more comprehensive framework for understanding how cAMP regulates fibrosis resolution across cell types, organs, and stages of tissue repair. An additional pertinent consideration relates to the relative expression of Gαs-coupled GPCRs and cAMP-specific phosphodiesterase isoforms within various cell types in different organs, as this will influence future drug development of cAMP-based therapies.

Our study has a number of limitations. One is the relatively low dose of bleo (1.0 mg/kg) used herein as compared to some other studies. Although this was intentional in light of our desire to investigate and characterize spontaneous resolution, it very likely limited the number of *Cthrc1^+^* fibroblasts as well as certain populations of aberrant epithelial cells. Indeed, the salutary impact of fibroblast cAMP in more severe and persistent models of fibrosis warrants exploration. Additionally, we did not enrich for particular cell types prior to performing scRNA-seq – unlike many other studies – thus reducing the resolution power to detect rare subtypes. Finally, as we did not employ scRNA-seq in our skin model, we cannot determine whether the types of changes in cellular subsets and intercellular crosstalk observed during resolution in the lung likewise apply to resolution in the skin. Nevertheless, our study establishes timely fibroblast-specific cAMP activation as a requirement for fibrosis resolution and associates its signaling with fibroblast apoptosis and pro-resolution cellular crosstalk. These findings emphasize and provide mechanistic context to the recently demonstrated clinical efficacy of the cAMP-elevating therapies nerandomilast (*77*) and treprostinil (*70, 71*).

## METHODS

### Sex as a biological variable

For animal experiments modeling lung fibrosis, we examined male and female animals, and no meaningful differences were noted between the sexes. For animal experiments modeling skin fibrosis, female cohorts were used given the female predominance of systemic sclerosis and to minimize the sex-related variability in baseline dermal thickness and fibrotic responses (*78*).

### Animal studies

WT male and female C57BL/6 mice were obtained from The Jackson Laboratory and used at 8–12 weeks of age. Transgenic mice containing *loxP* sites flanking exon 1 of the *Gnas* locus (The Jackson Laboratory, strain no. 035239) were crossed with *Col1a2*^CreERT2+/0^ mice (The Jackson Laboratory, strain no. 029567) to generate tamoxifen-inducible, fibroblast-specific conditional Gαs-KO mice. Genotyping was performed using genomic DNA extracted from tails. Briefly, genomic DNA was extracted using the REDExtract-N-Amp Tissue PCR Kit (Millipore Sigma). Cre genotyping was performed by PCR (PCR) using a *Col1a2-*Cre– specific primer pair (forward, 5′-CAGGAGGTTTCGACTAAGTTGG-3′ and reverse, 5′-CATGTCCATCAGGTTCTTGC-3′). PCR primers designed to bind to exon 5 and the 3′ *loxP* were used (5′-CGAGAGCGAGAGGAAGACA-3′ and 5′-GAGCTTCTGGCACATTCTTCT-3′, respectively) to verify the homozygosity of the floxed allele for *Gnas*. PCR primers designed to flank the *loxP* regions of the *Gnas^fl^* locus (forward, 5′-TGACCCCAAGACAGCACAG-3′ and reverse, 5′-GTGGCCTTGGCCTCTCTTT-3′) were used to confirm successful recombination following tamoxifen administration in Cre^+^ mice. Rosa26 tdTomato genotyping was confirmed by PCR using a tdTomato forward primer (5’-CTGTTCCTGTACGGCATGG-3’) paired with WPRE reverse primer (5’GGCATTAAAGCAGCGTATCC-3’).

Pulmonary fibrosis was induced in mouse lungs following single-dose o.p. bleo administration (1.0 U/kg, Millipore Sigma, B5507), as described previously (*20*). To determine the role of Gαs during spontaneous fibrosis resolution, its fibroblast-specific deletion was achieved through tamoxifen-induced Cre activation. Specifically, tamoxifen chow (40 mg/mouse/day; Inotiv, TD.130859) was administered starting on day 6 (to express tdTomato in controls) or day 21 (to delete Gαs during the resolution phase) and continued until harvest at the indicated time points. Mice were sacrificed on days 21, 42, or 63 to capture peak lung fibrosis and at mid- and late-resolution, respectively. Lungs were perfused with cold PBS, and the right upper, middle, and left lung lobes were harvested to assess fibrotic endpoints. Specifically, lung lobes were pooled in PBS, homogenized, and assessed for hydroxyproline content to estimate total collagen and whole-lung RNA, while the right lower lobe was used for fibroblast isolation and culture *in vitro*, or sectioned to perform Masson’s trichrome staining, followed by imaging with bright-field, whole-slide imaging with a Vectra Polaris Brightfield Scanner. The histologic severity of pulmonary fibrosis of Masson’s trichrome stained sections was assessed using the modified Ashcroft scale (*79*). Each sampled field was assigned a score from 0 (normal lung architecture) to 8 (complete obliteration of the field by fibrosis) based on the extent of alveolar septal thickening and disruption of lung architecture. For each lung, blinded scorers assessed at least 15 digital microscopic images (20x objective) per lung and the scores were averaged. The volume fraction of abnormal parenchymal epithelium was estimated by stereological point counting on histologic lung sections (*80*). A point grid was superimposed on each sampled field, and the percentage was calculated as 100 × the number of points falling on abnormal parenchymal epithelium divided by the total number of points falling on the respiratory zone (distal lung parenchyma, excluding terminal airways and adjacent vasculature). Morphometric measurements were made by a pathologist blinded to study group using ImageJ 1.8.0 (NIH, USA). Skin fibrosis was induced by daily subcutaneous injection of bleo (10 mg/kg, TEVA) divided over two regions on the flank for 7 days. For skin tissue readouts, punch biopsies of each injection area were obtained at the time of harvest (day 24 or 63) and snap frozen for hydroxyproline analysis or kept in 10% formalin for 72 h followed by 70% ethanol and paraffin embedding prior to Masson’s trichrome staining.

### Immunofluorescence microscopy

Mouse lungs were fixed with 10% formalin, embedded in paraffin, sectioned, and mounted on glass slides. Subsequent deparaffinization, citrate antigen retrieval, blocking and staining were performed as previously described (7). Staining of fixed fibroblasts and murine lung tissue was performed with the following primary antibodies: Cd63 (1:500, Sigma HPA010088), Arg1 (1:500, Proteintech 16001-1-AP), Trem2 (1:500, Proteintech, 68723-1-Ig) Cthrc1 (Proteintech, 16534-1-AP), Csmd1 (Bioss, BS-1579R), αSMA-Cy3 conjugated (Millipore Sigma, C6198), tdTomato (Rockland, 600-401-379), Ppdn (1:100, University of Iowa DSHB 8.1.1), Cldn4 (1:100, Invitrogen 364800), Krt8 (1:20, University of Iowa Developmental Studies Hybridoma Bank (DSHB) TROMA-I), Pdgfrα (LS Bio B2683), and p-Creb (Abcam, AB32096) overnight at 4°C. Unconjugated antibodies were then incubated with anti-rabbit secondary antibodies: FITC (1:200; Jackson 711-545-152), Cy3 (1:200, Thermo Fischer AlexaFluor 568), or Cy5 (1:200; Jackson 107-605-142) as indicated for macrophage and fibroblast staining. For epithelial staining, the following secondary antibodies were used: Cy5 (Anti-Syrian Hamster A647 1:100, Jackson Labs, 107-605-142) and Cy3 (anti-Rat 1:250, Jackson 712-166-150) as indicated. Mounting medium containing DAPI (Invitrogen) was used to stain nuclei. Images were obtained using a Nikon Eclipse Ti2 Inverted Confocal or an Olympus BX53 microscope.

### scRNA-seq data processing and analysis

Raw sequencing data were processed using CellRanger (version 7.2.0, 10x Genomics). Sequencing reads were aligned to the *Mus musculus* reference transcriptome (mm10) using the STAR aligner. The CellRanger count pipeline was run with default parameters to generate gene expression count matrices. Raw and filtered feature barcode matrices were imported into R (version 4.3.1) using the Read10X function. All downstream analysis was performed in R using Seurat (version 5.0.0). Four datasets, corresponding to distinct experimental time points (saline control, day 21, day 42, day 63) were preprocessed independently. Each dataset consisted of high-quality singlet cells that underwent quality control as described below. Ambient RNA contamination was corrected using SoupX (version 1.6.2). A SoupChannel object was created from the raw and filtered matrices, and the contamination fraction was estimated from the expression profiles of empty droplets. This estimated background was then used to adjust expression counts for individual cells, and corrected count matrices were saved for downstream analysis. Standard quality control thresholds were applied to remove low-quality cells and potential multiplets (Supplemental Figure 3B). Cells with fewer than 200 detected genes or more than 10,000 genes, fewer than 500 or more than 20,000 total RNA counts, or greater than 10% of UMIs mapping to mitochondrial genes were excluded. Filtered data were normalized using log-normalization, regressing out total UMI counts and the percentage of mitochondrial transcripts per cell. Variable features were identified using the VST method, and principal component analysis (PCA) dimensionality reduction was performed. Data were further normalized using SCTransform (glmGamPoi method) to stabilize variance across sequencing depth. The first 30 principal components were used for UMAP embedding and unsupervised clustering based on a shared nearest neighbor with a resolution of 0.5. Doublet detection was performed using DoubletFinder (version 2.0.4). The expected doublet rate was set to 7.5% based on 10x Genomics guidelines for the estimated cell recovery rate (*20*), and default parameters were used otherwise. Cells classified as singlets were then extracted for downstream analysis.

Cell type annotation was conducted with Clustifyr (version 1.8.0). Cluster-level expression profiles were correlated to two mouse lung single-cell reference datasets from CZ CellxGene and ScTypeDB (21, 22). Gene identifiers were harmonized to *Mus musculus* gene symbols using biomaRt. Cluster-level correlations were computed using clustify and cell type identities were assigned to each cluster. Manual review and refinement of annotations was performed as needed. Differential gene expression analysis was conducted across annotated cell types to identify key marker genes. Subsets of selected cell populations (fibroblasts, epithelial cells, and macrophages) were reanalyzed using the same Seurat workflow, with clustering resolution increased to 0.7 to capture finer substructure within lineages. Investigation of cell-cell communication networks was performed using CellChat (version 1.6.1) on the datasets. Analyses were performed both on the full dataset and on a subset of selected cell types using metadata from the Seurat cluster identities from the SCT assay. For presentation and clarity, representative terms were selected from related groups of GSEA gene sets to reduce redundancy. The CellChat mouse ligand-receptor database (including non-protein interactions) was used to predict signaling interactions, with cell populations with fewer than 10 cells excluded from the analysis. Overexpressed genes and ligand-receptor interactions were identified, followed by computation of communication probabilities between cell types and pathway and centrality analyses. All CellChat analyses were run with default parameters for functions and visualization. Gene module scores were calculated as previously described (*20*).

### Reagents

Unless otherwise specified, pharmacologic agents were reconstituted in DMSO as stock solutions and stored at –80°C with working concentrations indicated in parentheses. Recombinant human TGFβ (2 ng/mL for human fibroblasts, 5 ng/mL for mouse fibroblasts) was purchased from R&D (7754-BH) and resuspended in filter-sterilized 1% BSA. PGE_2_ (500 nM for human fibroblasts, 250 nM for mouse fibroblasts) and the direct adenylyl cyclase activator forskolin (20 µM) were purchased from Cayman Chemicals (14010, 11018).

### Cells

MRC5 normal fetal human lung fibroblasts were obtained from the American Type Culture Collection. All cells were initially cultured in low glucose DMEM (Invitrogen) supplemented with 10% FBS (Hyclone), 100 units/mL penicillin, and 100 μg/mL streptomycin (both from Invitrogen). Cells were then serum starved in FBS-free DMEM overnight, and differentiation to MFs was induced by treatment with TGFβ for 48 h. TGFβ–elicited MFs were then treated for specified time points as described followed by harvesting. Fibroblasts from lung tissue of saline- or bleo-treated mouse lungs were obtained during lung harvest at the indicated times as previously described (*20*). For all *in vitro* experiments, a minimum of three independent experiments were performed, and the results are presented as mean ± SEM.

### CRISPR/Cas9 studies

Stable inducible fibroblast lines capable of CRISPR/Cas9-mediated deletion of Gαs were generated via infection with lentiviral particles containing pLentiCRISPR v2 TLCV2 plasmids (Addgene no. 87360) controlling the following sgRNA sequences: *GNAS*, 5′-CACCGGTCGGGGTGTAGCTTACGGG-3′. The NT sequence 5′-AAATGTGAGATCAGAGTAAT-3′ (Thermo Fisher Scientific, A35526) was likewise incorporated into TLCV2 and used as a negative control. Polybrene (2 μg/mL, Millipore Sigma, TR-1003-G) was added to lentiviral suspensions to enhance transduction efficiency in MRC5 cells followed by puromycin selection (5 μg/mL) for 48–72 hours. Induction of CRISPR/Cas9-mediated deletion of Gnas was achieved by addition of doxycycline (1 μg/mL, Cayman Chemicals) to conditioned medium to promote expression of the *GNAS*-specific sgRNA.

### Western blot analysis

Cells were lysed in RIPA buffer supplemented with protease inhibitors (Roche Diagnostics, 11836153001) and a phosphatase inhibitor cocktail (EMD Biosciences, 524624 and 524625). Proteins were separated by SDS-PAGE and transferred to a nitrocellulose or polyvinylidene difluoride membrane. Membranes were subsequently blocked with 5% BSA and probed with a mouse antibody specific to Gαs (Abcam, AB97629), αSMA (Agilent, M0851), collagen 1A1 (CST, 91144), P-CREB S133 (Abcam, AB32096) and GAPDH HRP conjugate (CST, 8884). GAPDH was used as a loading control for all remaining Western blots.

### Apoptosis assays

Apoptosis was determined either by extracellular Annexin V expression or by TUNEL staining of fixed lung sections (Roche, 11684795910). For Annexin V readouts, fibroblasts or MFs were treated with 100 ng/mL of anti-Fas activating antibody (EMD Biosciences, CH11 05-201) overnight to induce apoptosis. Annexin V+ cells were quantified using flow cytometry (BD Fortessa flow cytometer). TUNEL staining of mouse lung tissue was achieved using citrate antigen retrieval for 8 min followed by signal development per the manufacturer’s protocol.

### Statistics

Statistical analysis was performed using GraphPad Prism, version 11.0.0 (GraphPad Software). Experimental data are presented as the mean and were analyzed for statistical significance by 1-way ANOVA with Tukey’s multiple-comparison test or paired/unpaired 2-tailed *t* test, as appropriate. A *P* value of less than 0.05 was considered significant. Data represent the mean ± SEM.

### Study approval

All animal experiments were approved by the University of Michigan Institutional Animal Care and Use Committee (IACUC) and are reported in accordance with the Animal Research: Reporting of In Vivo Experiments (ARRIVE) guidelines.

### Data availability

The scRNA-seq datasets generated in this study are publicly available in the Gene Expression Omnibus (GEO) under accession number GSE328925. All other data supporting the findings of this study are available from the corresponding author upon reasonable request.

## Acknowledgments

This work was funded by NIH grants K08 HL163178 (to SMF), R01 HL175555 (to TB), Hevolution HF-AGE award (to C.A.A.), National Science Foundation CAREER award 2045977 and 2548872 (to CAA), the Department of Defense and Congressionally Directed Medical Research Program W81XWH2010336 and W81XWH2110491 (to CAA), Defense Advanced Research Projects Agency (DARPA) “BETR” award D20AC0002 (to CAA) awarded by the U.S. Department of the Interior (DOI), Interior Business Center, National Institute of Health. R01 AR074997 (to JV), R01 AR087567 to CAA, R35 HL160770 (to RLZ), R35 HL144979 (to MPG). All Figure schematics were created with BioRender (BioRender.com).

## Author contributions

SMF, NMW, and MPG designed the experiments. Experiments were performed by SMF, JS, JDB, KW. Data were analyzed by SMF, JS, NMW, VTW, SDG, RH, VP, HP, HR, CAA, RLZ, and MPG. Bioinformatics were performed by JS, VTW, SDG, and YL. Patient-derived normal and IPF fibroblasts and lung sections were provided by SKH. Study was conceptualized by SMF and MPG with additional intellectual contributions provided by TSB, JV, SKH, CAA, and RLZ. The manuscript was written by SMF and MPG. All authors reviewed the manuscript.

## Supplemental Figures

**Supplemental Figure 1.**
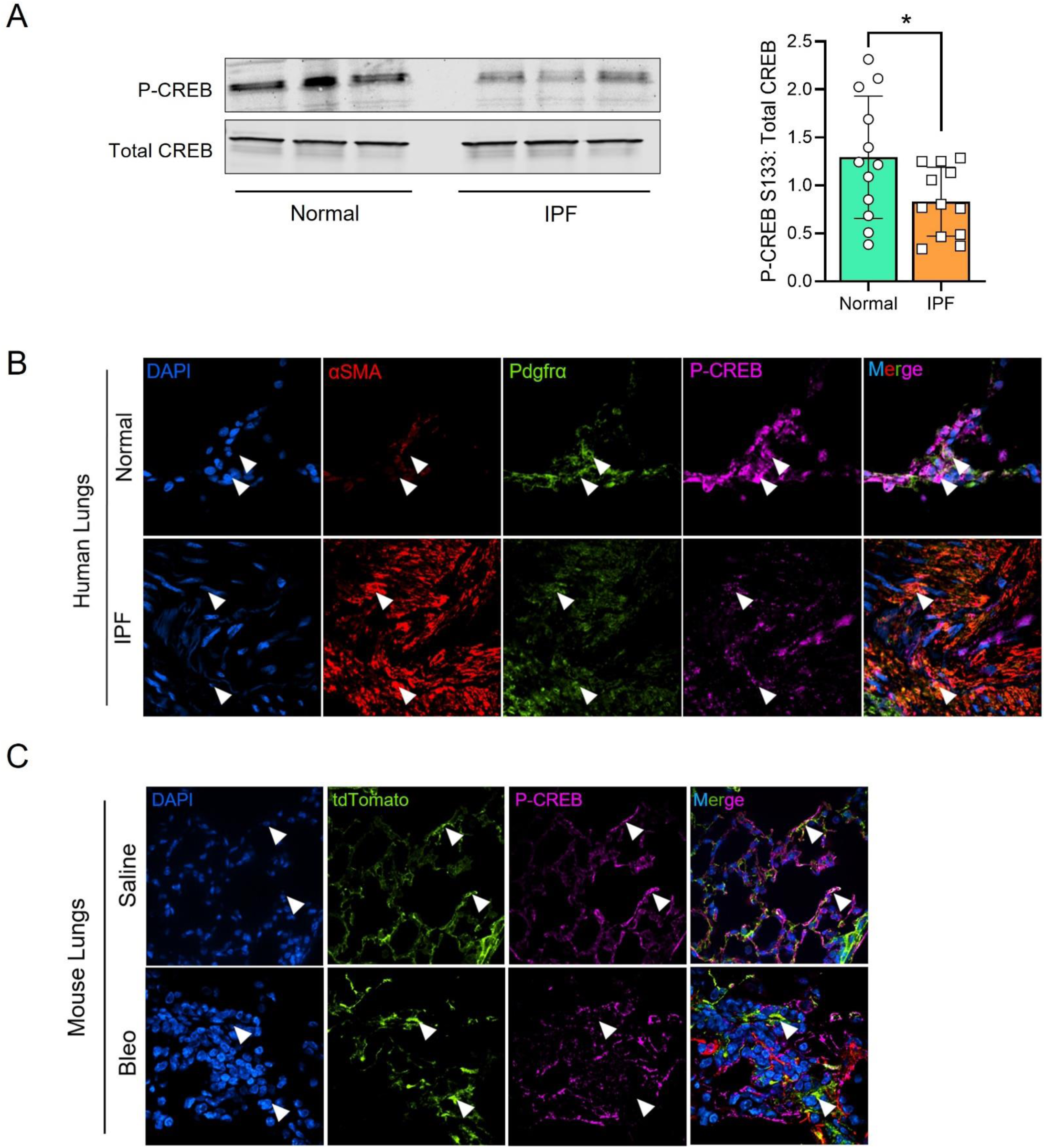
(**A**) Quantification of P-CREB protein by Western blot in fibroblasts derived from the lungs of normal healthy and IPF patients (left: representative blots, right: densitometric analysis). A total of 12 distinct healthy and IPF lines were used indicated by the number of data points in each histogram. \**P <* 0.05 as determined by 2-tailed t-test. (**B**) Immunofluorescence staining of P-CREB, αSMA, and PDGFRα in normal healthy human lungs and IPF lungs. (**C**) Immunofluorescence staining of P-CREB and tdTomato in saline- and bleo-treated mouse lungs at day 21.

**Supplemental Figure 2.**
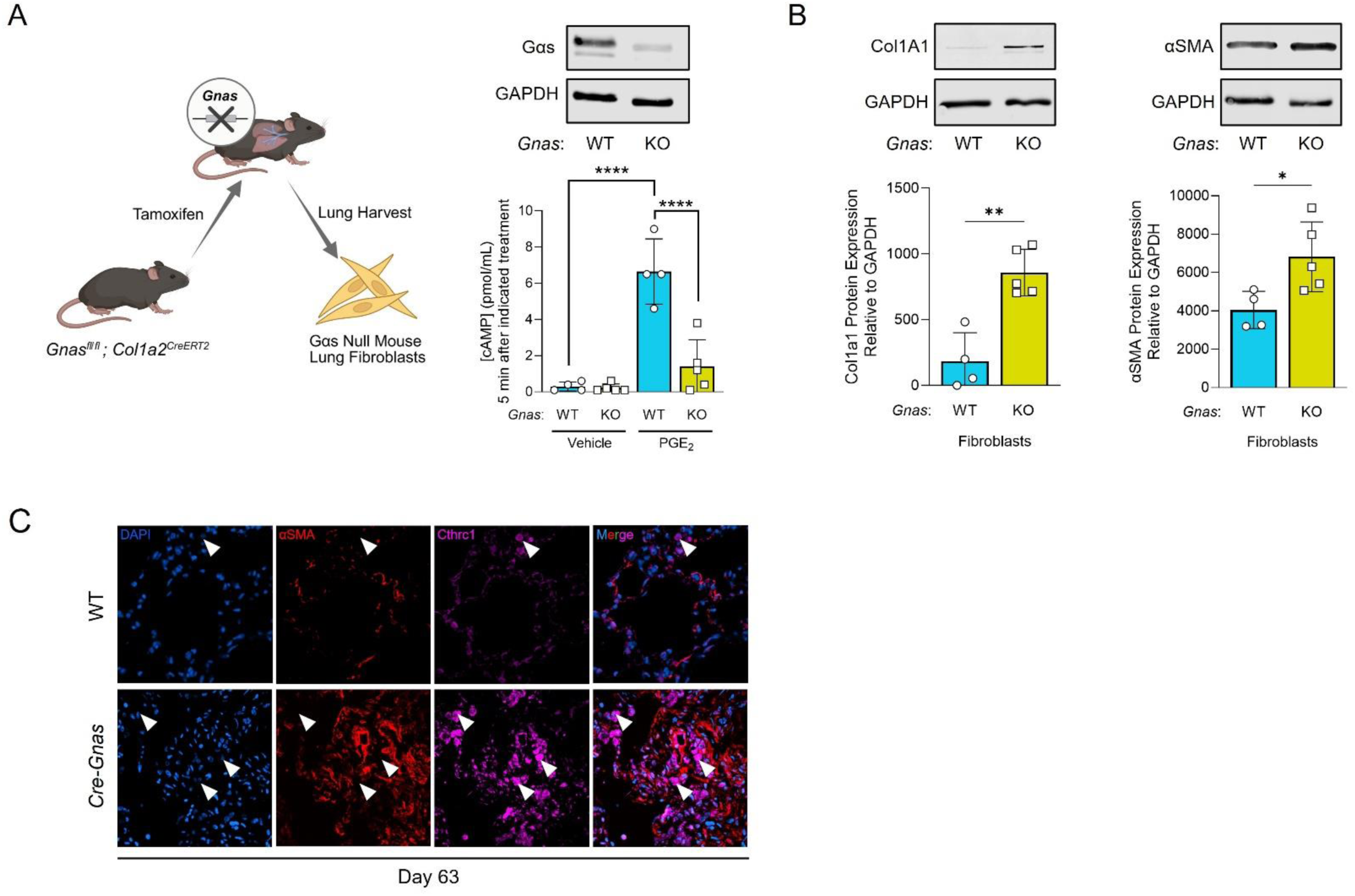
(**A**) Fibroblast-specific Cre-mediated *Gnas* deletion (schematic, left), confirmation of Gαs protein deletion (top right), and quantification of intracellular cAMP-generation following PGE_2_ treatment in fibroblasts derived from WT and *Col1a2Cre*;*Gnas* KO mice. (**B**) To ensure a pure fibroblast culture, cells were passaged 4 times prior to protein quantification by Western blot of fibrosis-associated genes Col1A1 and αSMA in unstimulated fibroblasts maintained in culture with serum free medium for 48 h (top: representative blots, bottom: densitometric analysis). (**C**) Representative immunofluorescence staining of αSMA and Cthrc1 in day 63 WT and *Col1a2Cre*;*Gnas* KO lung sections.

**Supplemental Figure 3.**
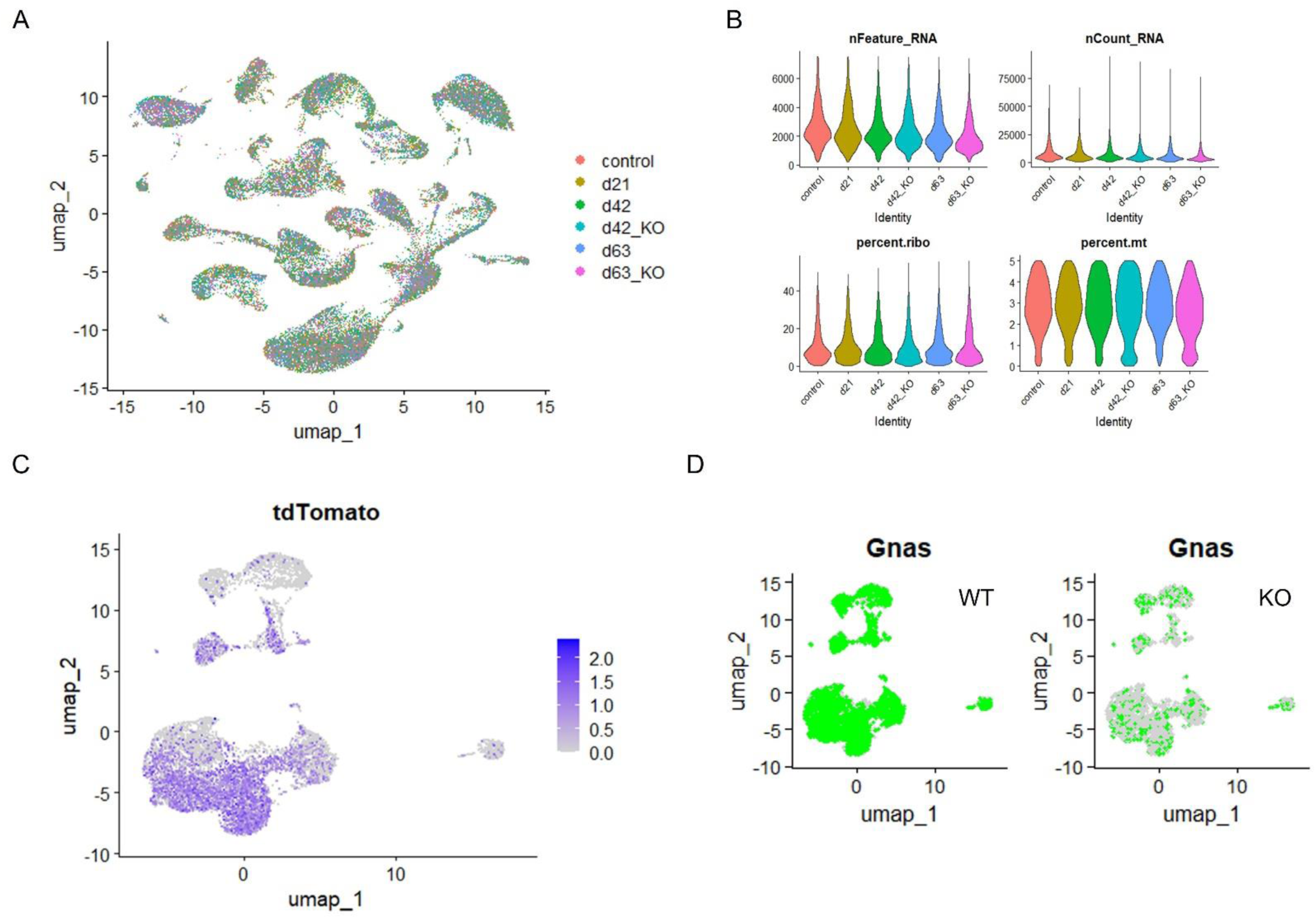
(**A**) Superimposed UMAP of all conditions and time points. (**B**) Standard quality control thresholds for unique genes/cell (upper left), unique molecular identifiers/cell (upper right), ribosomal percentage (lower left), and mitochondrial percentage (lower right). (**C**) Mesenchymal cell subclustered UMAP feature plot of overall tdTomato expression. (**D**) Mesenchymal cell subclustered UMAP feature plot of *Gnas* expression in WTs (left) versus *Col1a2;Gnas* KOs (right).

**Supplemental Figure 4.**
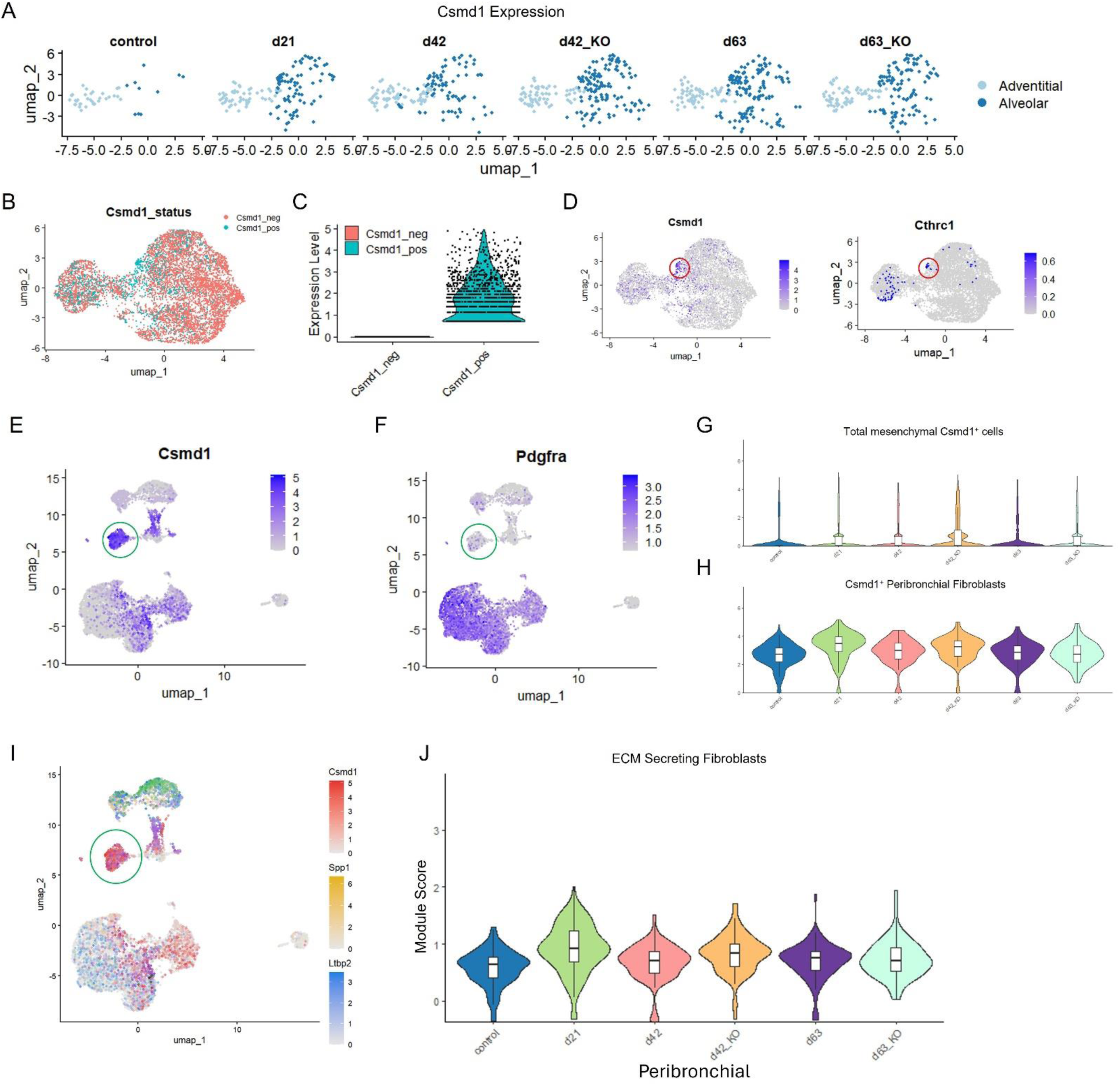
(**A**) Longitudinal display of *Csmd1* expressing cells among adventitial (light blue) and alveolar fibroblasts (dark blue) in control and bleo-treated as well as WT and *Col1a2;Gnas* KO mice. (**B**) Feature plot highlighting *Csmd1*^-^ and *Csmd1*^+^ cells within adventitial and alveolar fibroblast subclustered UMAP. (**C**) Quantification of Csmd1 expression level in *Csmd1*^-^ and *Csmd1*^+^ clusters. (**D**) Feature plot displaying relative *Csmd1* (left) and *Cthrc1* (right) expression levels within adventitial and alveolar fibroblasts. (**E**) Mesenchymal cell subclustered UMAP feature plot of *Csdm1* and *Pdgfra* (**F**). (**G** & **H**) Violin plots displaying total number of *Csdm1* positive cells at each time point and condition among all mesenchymal cells (**G**) and peribronchial fibroblasts (**H**). (**I**) Feature plot displaying ECM-secreting fibroblast signature genes. (**J**) Violin plot of module scores assigned to *Csmd1*, *Spp1*, and *Ltbp2* expression. The red circles in **D** encompass a dense region of *Csmd1*^+^ and *Cthrc1*^+^ fibroblasts. The green circles in E, F, and I identify the peribronchial fibroblast subpopulation.

**Supplemental Figure 5.**
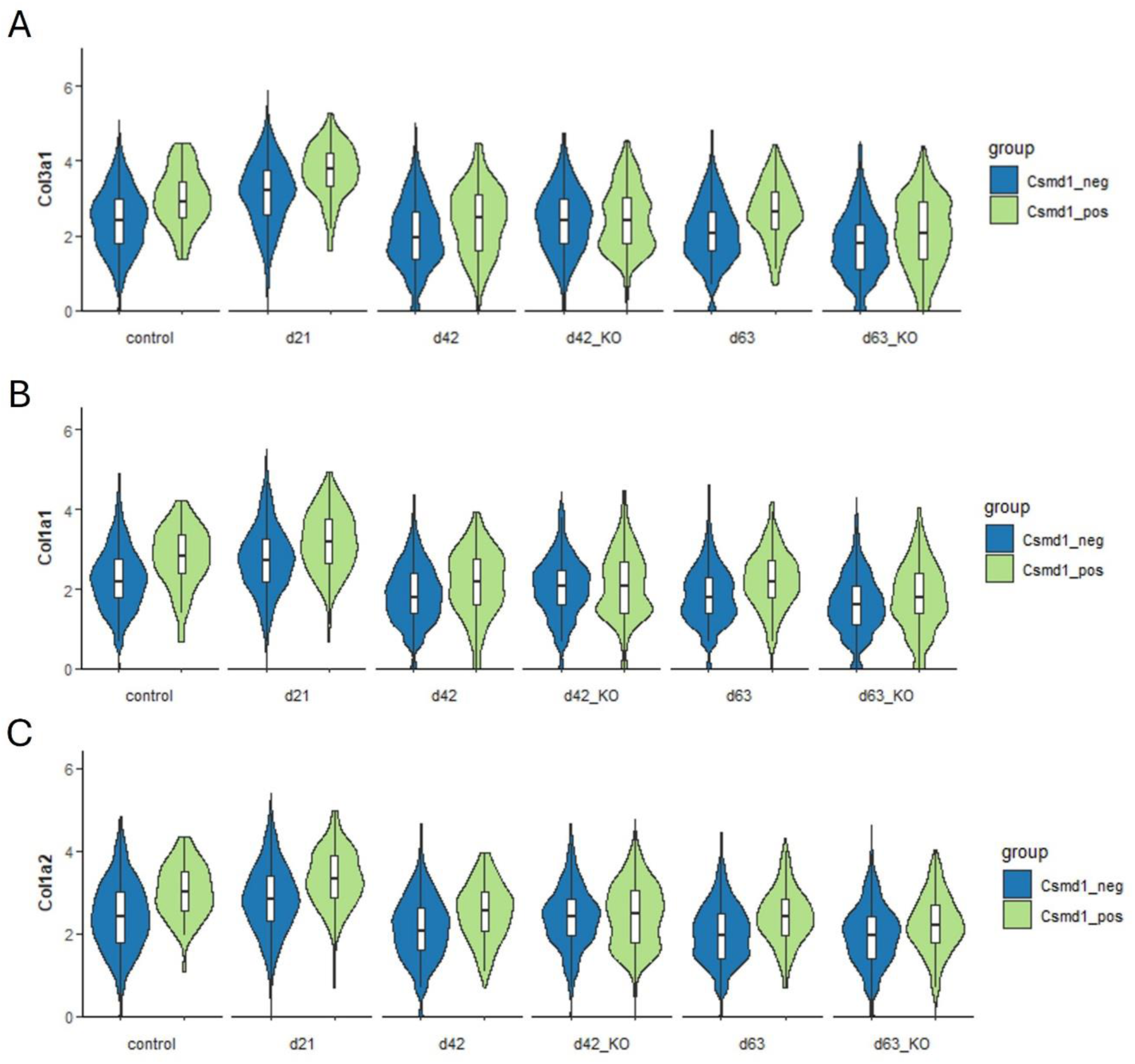
(**A-C**) Violin plots displaying relative expression of the collagen genes *Col3a1* (**A**), *Col1a1* (**B**), and *Col1a2* (**C**) among *Csmd1*^-^ and *Csmd1*^+^ cells at each time point and condition.

**Supplemental Figure 6.**
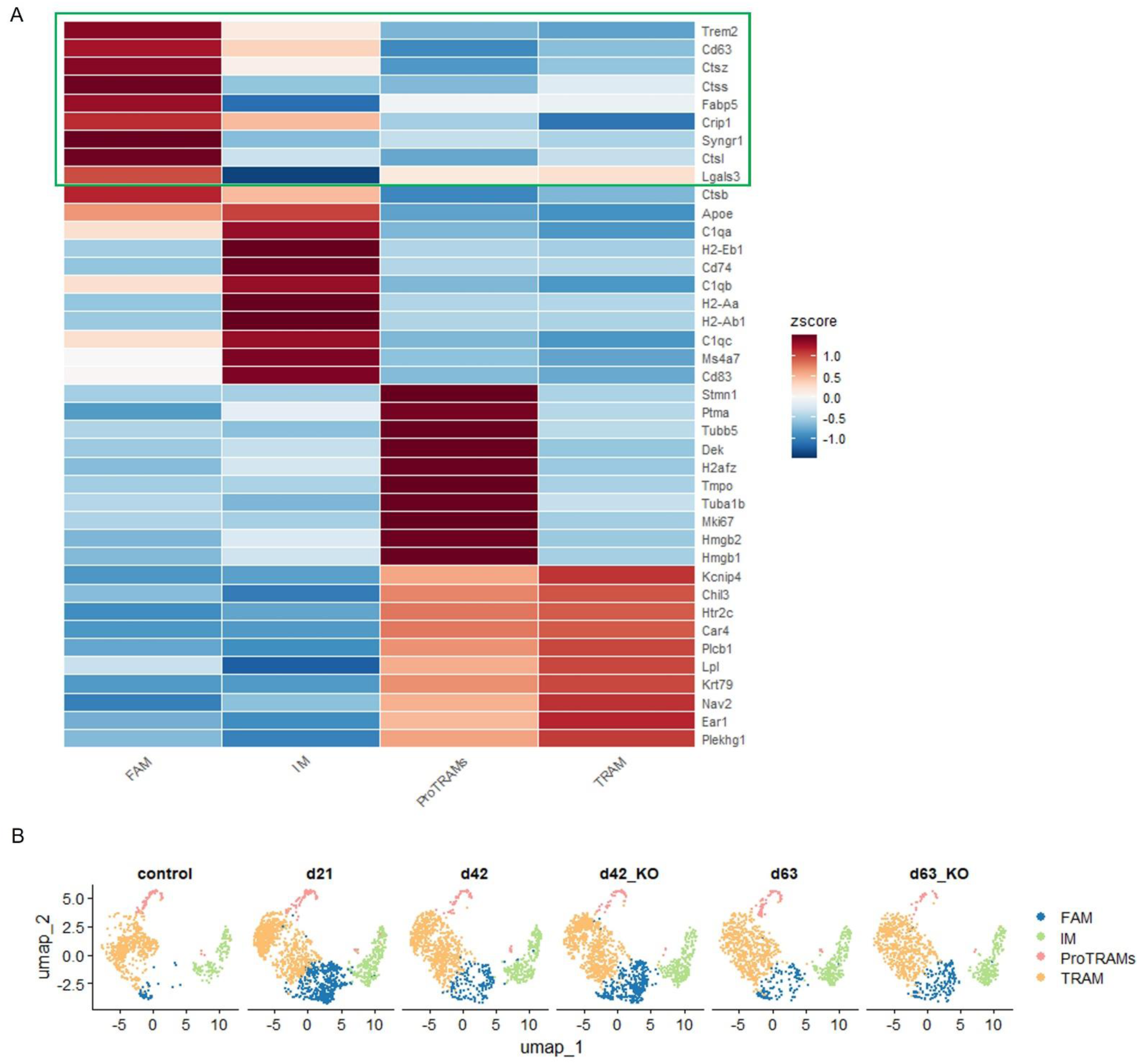
(**A**) Heatmap of the highest expressed genes by lung macrophage subtype. The green box outlines the top genes expressed in FAMs. (**B**) Macrophage subclustered UMAPs displaying relative proportions of FAMs, IMs, proTRAMs, and TRAMs at each time point and condition.

**Supplemental Figure 7.**
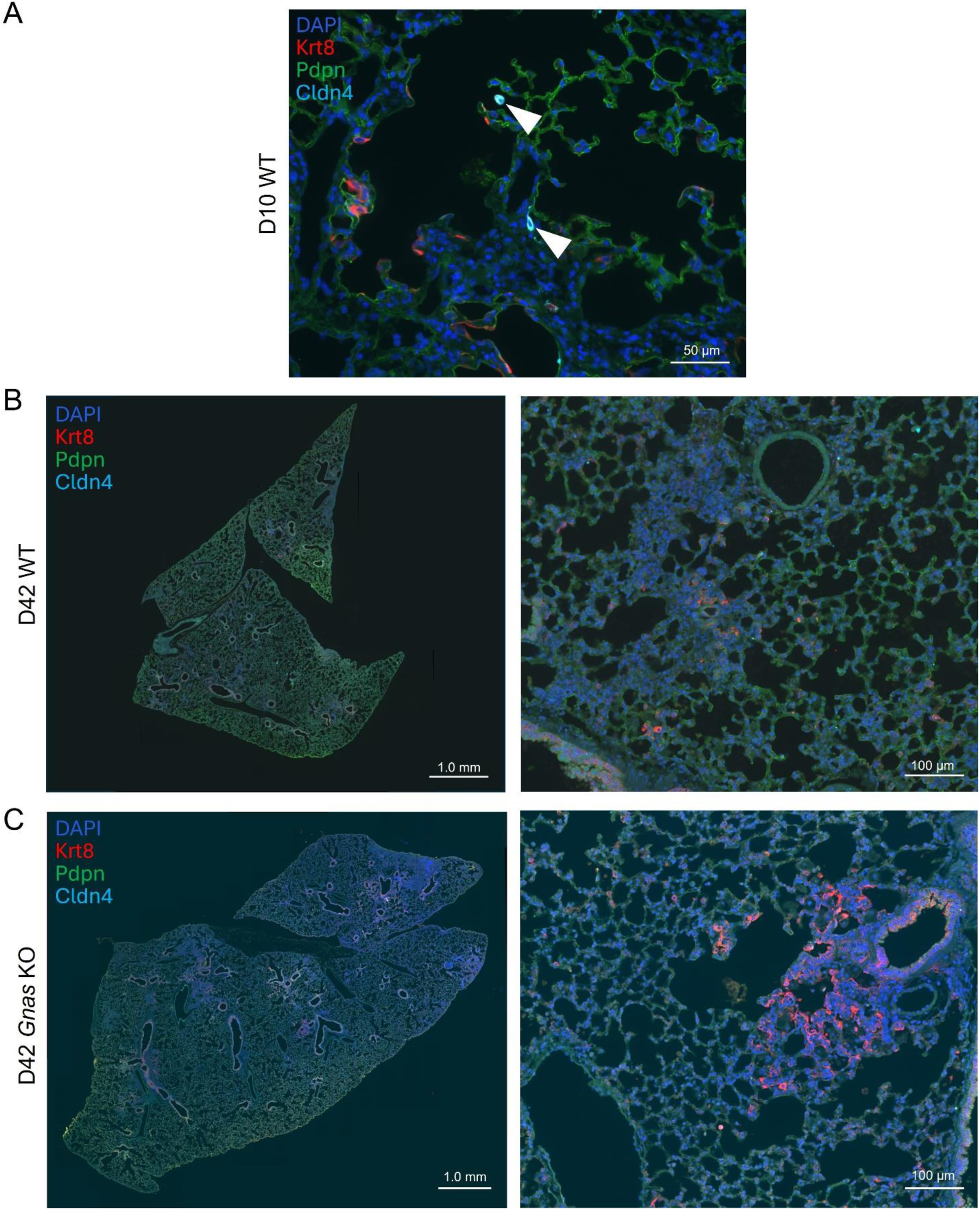
Immunofluorescence staining for Cldn4, Krt8, and Pdpn day 10 (positive control) (**A**), day 42 post-bleo WT (**B**), and day 42 *Gnas* KO (**C**) post-bleo. White arrows in (A) show Cldn4 positive cells.

**Supplemental Figure 8.**
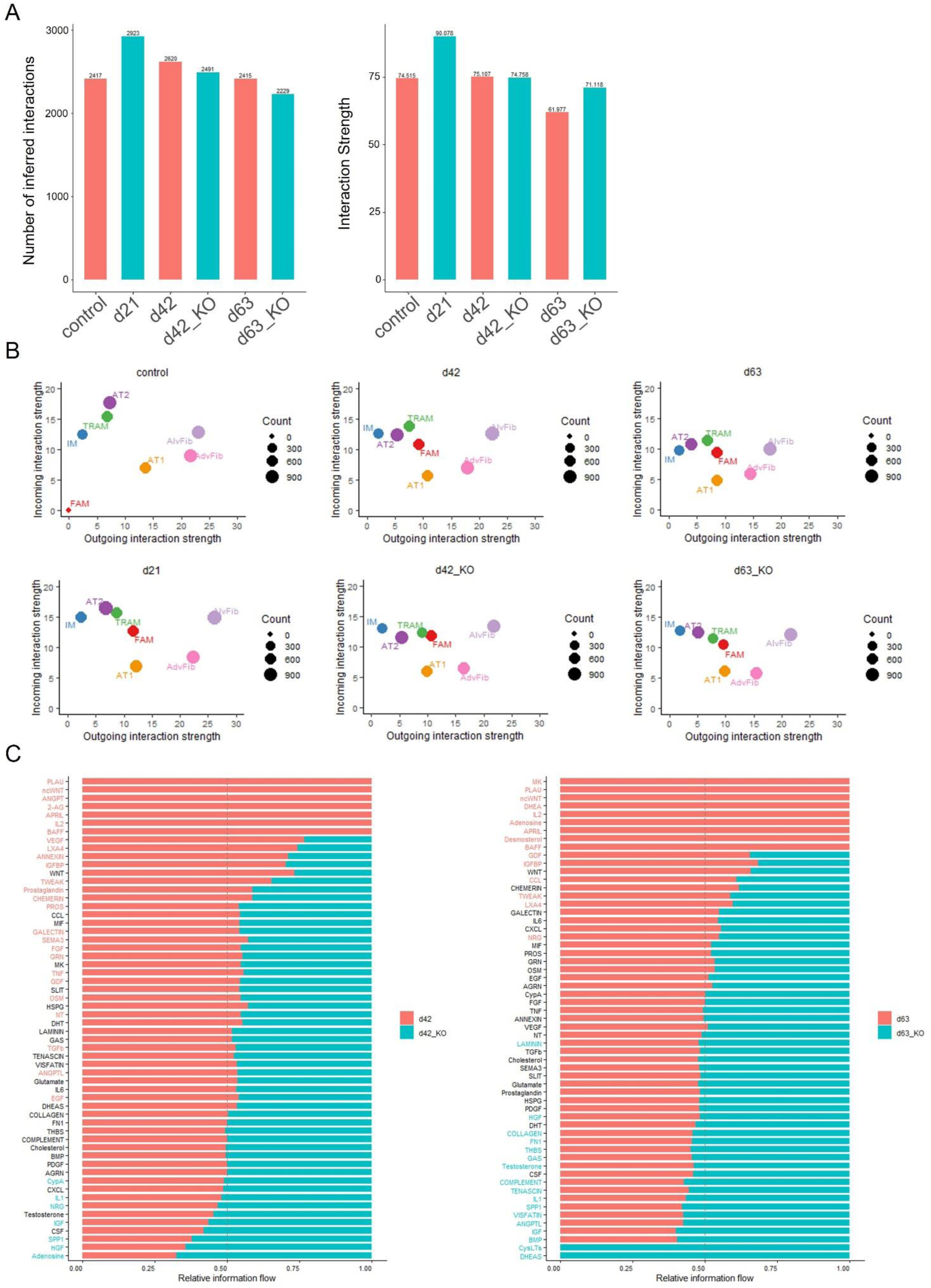
(**A**) Histogram displaying total number of inferred interactions (left) and interaction strength (right) at each time point and condition. (**B**) Dot plot depicting relative incoming and outgoing interaction strength among the indicated cell types at each time point and condition. (**C**) Waterfall plots displaying relative enrichment of the indicated pathways. Each plot compares WT and KO on days 42 (left) and 63 (right).

**Supplemental Figure 9.**
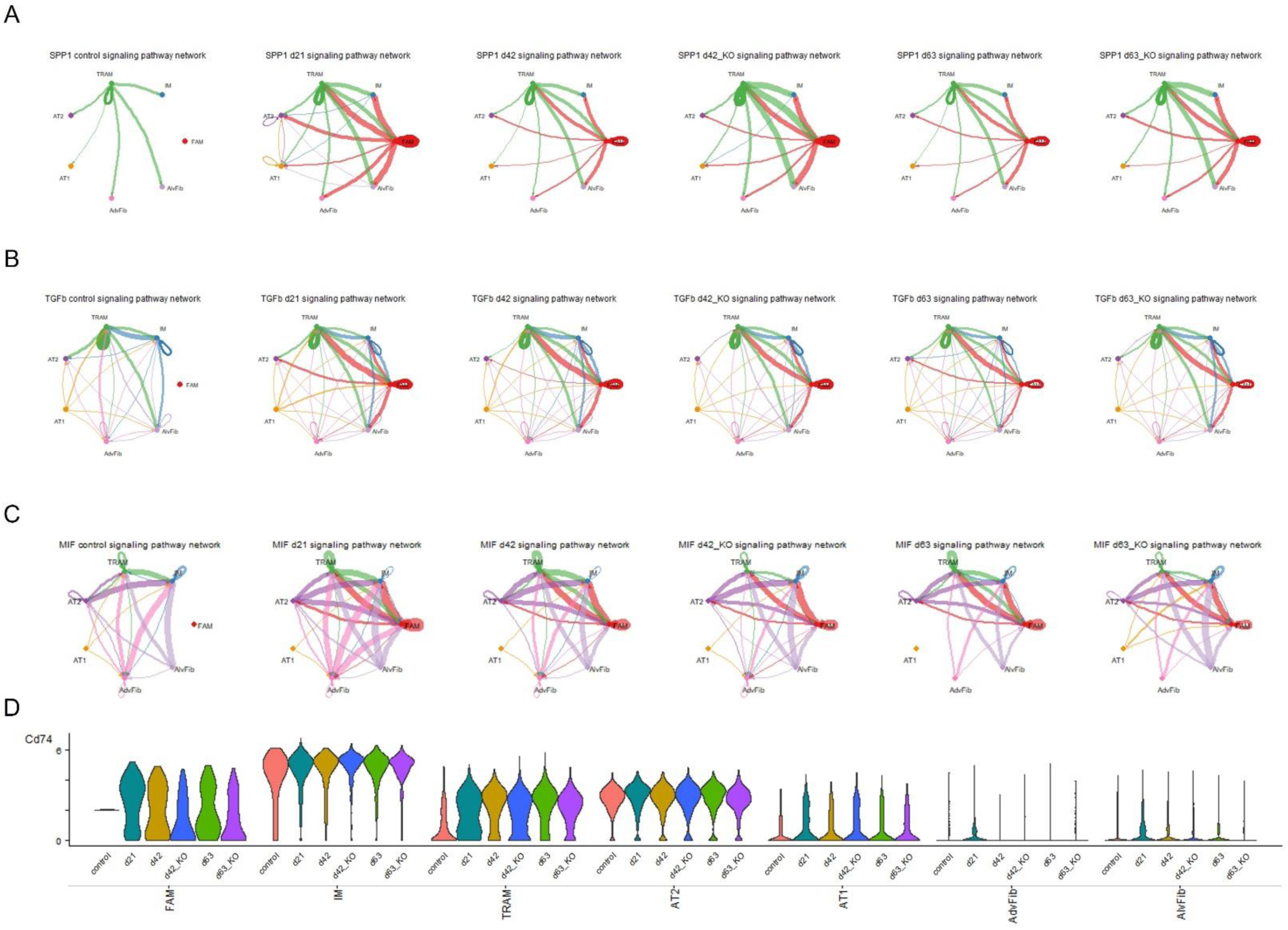
(**A**–**C**) Circos plots displaying cellular crosstalk of the SPP1 (**A**), TGFβ (**B**), and MIF (**C**) pathways at each time point and condition. Line thickness is proportional to signaling strength, line color indicates sender cell. (**D**) Violin plots displaying relative expression of the MIF receptor gene *Cd74* in the indicated cell subtype at each time point and condition.

**Supplemental Figure 10.**
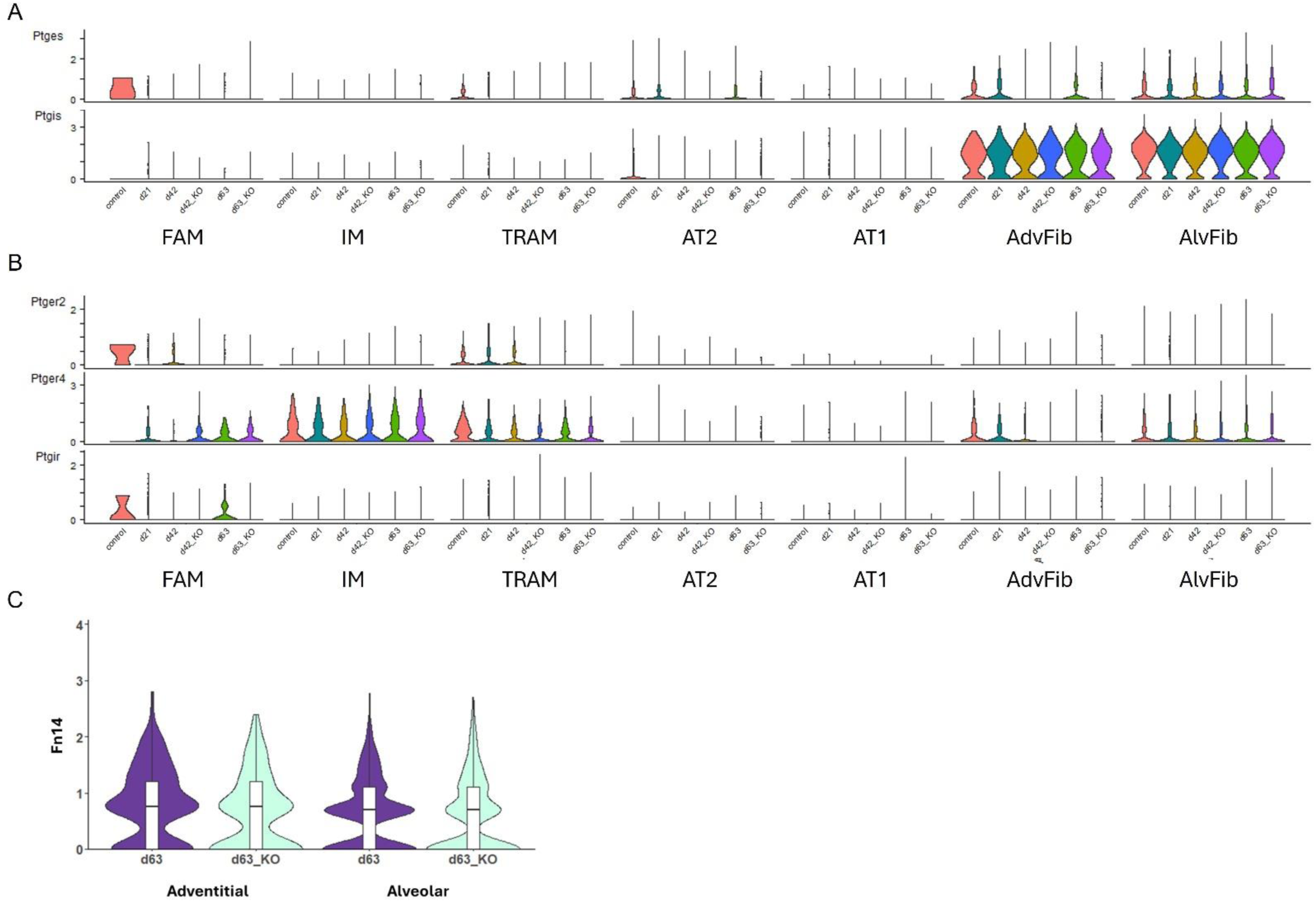
(**A**) Violin plots displaying relative expression of prostaglandin synthase genes for PGE_2_, (*Ptges*, top) and PGI_2_ (*Ptgis*, bottom) in the indicated cell subtype at each time point and condition. (**B**) Violin plots displaying relative expression of prostaglandin ligand receptor genes for the PGE_2_ receptors EP2 (*Ptger2*, Top), EP4 (*Ptger4*, second from top) and PGI_2_ receptor (Ptgir, bottom) in the indicated cell subtype at each time point and condition. (**C**) Violin plots displaying relative expression of the TWEAK receptor gene *Fn14* within adventitial and alveolar WT and KO fibroblasts on day 63.

